# Deconstructing the role of iron and heme in the structure of healthy human gut microbial communities

**DOI:** 10.1101/2022.01.25.477750

**Authors:** Arianna I. Celis, David A. Relman, Kerwyn Casey Huang

**Affiliations:** Department of Bioengineering, Stanford University, Stanford, CA 94305; Department of Medicine, Stanford University School of Medicine, Stanford, CA 94305; Department of Microbiology and Immunology, Stanford University School of Medicine, Stanford, CA 94305; Infectious Diseases Section, Veterans Affairs Palo Alto Health Care System, Palo Alto, CA, 94304; Chan Zuckerberg Biohub, San Francisco, CA 94158

## Abstract

The response of the human gut microbiome to disruptions is often difficult to discern without model systems that remove the complexity of the host environment. Fluctuations in iron availability provide a case in point: the responses of pathogenic bacteria to iron *in vitro* are much better understood than those of the indigenous human gut commensal microbiota. In a clinical study of iron supplementation in healthy humans, we identified gradual, participant-specific microbiota shifts correlated with bacterial iron internalization. To identify direct effects due to taxon-specific iron sensitivity, we used stool samples from these participants as inocula to derive *in vitro* communities. Iron supplementation of these communities caused small shifts in structure, similar to *in vivo*, whereas iron deprivation dramatically inhibited growth with irreversible and cumulative reduction in diversity and replacement of some dominant species. Sensitivity of individual species to iron deprivation during growth in axenic culture generally predicted iron dependency in a community context. Finally, exogenous heme acted as a source of inorganic iron to prevent depletion of some community members. Our results highlight the complementarity of *in vivo* and *in vitro* studies in deconstructing how environmental factors affect gut microbiome structure.

## Introduction

The gut microbiota is often subjected to environmental perturbations with long-term effects, including dietary shifts (Faith et al., 2011), osmotic diarrhea (Tropini et al., 2018), and antibiotics (Dethlefsen and Relman, 2011; Ng et al., 2019). Typically, the complexities of a mammalian host pose challenges for disentangling possible direct effects of the perturbation on the microbiota versus indirect effects of an altered host environment. Recent studies have shown that *in vitro* microbial communities can be used to model the gut microbiota’s response to antibiotics (Aranda-Díaz et al., 2020), other drugs (Javdan et al., 2020), and colonization resistance (Aranda-Díaz *et al*., 2020; Hromada et al., 2021), but it remains unclear to what extent *in vitro* communities can model other conditions, particularly ones that have especially profound effects on host cells.

A case in point is iron, which is essential for virtually all cells. It is incorporated directly into proteins involved in diverse processes such as DNA methylation and antibiotic biosynthesis (Andrews et al., 2003), and is a component of essential cofactors such as iron-sulfur clusters and heme that are involved in important functions such as ATP production, protection against reactive oxygen species, and oxygen transport (Andrews, 1998; Andrews *et al*., 2003; Gruss et al., 2012). Demand for iron in humans is met using dietary sources or supplements in the form of inorganic iron (Fe^2+^ or Fe^3+^) or heme-associated iron (Armitage and Moretti, 2019; Celis and Relman, 2020). Importantly, only 10-20% of ingested iron is absorbed in the small intestine (Armitage and Moretti, 2019; Kortman et al., 2014; Yilmaz and Li, 2018). The remainder travels to the colon, where the dense, colonic microbiota resides. Despite the essentiality of iron, high concentrations can also be toxic to bacteria. Hence, regulation of intracellular iron concentration is important, for both humans and the diverse set of bacteria within the gastrointestinal tract (Andrews, 2000; Andrews *et al*., 2003). Extracellular iron concentrations in the large intestine can fluctuate as a response to dietary and supplemental iron intake, absorption in the small intestine, and sequestration by the immune system, during times of inflammation, infection, or anemia (Kortman *et al*., 2014; Lund et al., 1999; Yilmaz and Li, 2018). Thus, it is important to determine the effects of changes to iron concentration on the gut microbiota in a community context and to establish the species-specific iron needs, acquisition strategies, and adaptation mechanisms of commensal gut bacteria.

Previous studies conducted on anemic infants living in low- and middle-income countries (LMIC) (Jaeggi et al., 2015; Zimmermann et al., 2010) and humans suffering from inflammatory bowel disease (IBD) (Kumar and Brookes, 2020) found that oral iron supplementation is associated with an increase in inflammatory markers (Jaeggi *et al*., 2015; Zimmermann *et al*., 2010), an increase in the relative abundance of pathogenic Enterobacteriaceae (Jaeggi *et al*., 2015; Zimmermann *et al*., 2010) and a decrease in the relative abundance of health-associated commensal bifidobacteria and lactobacilli (Jaeggi *et al*., 2015; Paganini and Zimmermann, 2017; Zimmermann *et al*., 2010). Such differential responses across taxa are perhaps to be expected, as the iron concentrations required for optimal growth of various bacterial species can differ by ∼100-fold (Andrews, 1998; Andrews *et al*., 2003). Pathogens such as *Escherichia coli* strains with extraintestinal virulence depend on iron for respiration and expression of virulence factors (Galardini et al., 2020). They produce high-affinity iron-binding molecules (siderophores) to scavenge iron when its concentration is low and hence compete with other bacterial species and the host for this essential metal. Furthermore, they are voracious when iron levels are high, storing extra iron atoms in their cytoplasm (iron content in *E. coli* is ∼10^5^ atoms/cell) for use during future conditions of iron deprivation (Abdul-Tehrani et al., 1999; Andrews *et al*., 2003; Lopez and Skaar, 2018). Pathogens can also acquire iron in the form of heme, which can be directly incorporated into heme-binding proteins or can be enzymatically degraded to release inorganic iron (Andrews *et al*., 2003; Cescau et al., 2007). By contrast to pathogenic *E. coli*, bifidobacteria and lactobacilli do not require iron for energy generation, do not produce siderophores, and contain as few as 1-2 iron atoms/cell (Andrews, 1998; Andrews *et al*., 2003). These differences are consistent with the growth advantages of pathogenic bacteria relative to commensals when iron levels are high.

Studies of iron deprivation in conventional mice have documented an associated drop in community diversity, in which members of the Prevotellaceae, Porphyromonadaceae, and Bacteroidaceae families appeared to go extinct (Coe et al., 2021). Studies conducted with *in vitro* fermentation models used the iron chelator 2,2’- dipyridyl to reduce iron concentrations available to a community derived from a malnourished child from an LMIC, and found a decrease in the relative abundances of the Lachnospiraceae and Bacteroidaceae families (Dostal et al., 2013; Dostal et al., 2015). 2,2’-dipyridyl is cell permeable and hence affects both extracellular and intracellular concentrations; thus, it is not ideal for representing real-world biological conditions.

The paucity of studies in which extracellular iron is controllably depleted means that the bacterial taxa within the highly diverse human microbiota most sensitive to iron deprivation in the absence of confounding disease-associated variables remain to be identified.

Here, we examined the effects of iron supplementation in healthy humans and then manipulated iron availability in communities derived *in vitro* using stool collected from these same humans to probe the iron requirements of commensal bacteria and the role of iron in shaping microbial community structure. In doing so, we sought to identify taxa that are resilient to changes in iron concentration and iron sources (i.e., inorganic iron versus heme-iron), in the absence of confounding host variables such as inflammation. We found that iron supplementation induces microbiota changes that are small, participant-specific, and positively correlated with increased bacterial intracellular iron concentrations during supplementation. We identified the Lachnospiraceae and Bacteroidaceae families as the most sensitive to iron supplementation and found that the specific genera and species affected within these families were participant-specific. Elevated iron availability in stable, diverse *in vitro* bacterial communities derived from our human participants recapitulated the moderate effects of iron supplementation *in vivo*, in contrast to iron deprivation, which induced irreversible changes to the community richness and composition that gradually accumulated. We found that resilience to iron deprivation varies across taxonomic families and that the variability of resilience to low iron within these families drives community shifts, as more resilient species occupy niches made available by less resilient ones. We discovered that the presence of heme buffers some of the effects of inorganic iron deprivation by rescuing specific species from depletion, and identify species that use heme as an alternate source of iron. Our results underscore the role of iron and heme availability in gut bacterial community structure, and highlight the general utility of *in vitro* communities for understanding the origins of human microbiota responses observed *in vivo*.

## Results

### Iron supplementation elicits participant-specific changes to gut microbiota composition in healthy humans

Inorganic iron supplements are available over-the-counter and are standard treatment for people diagnosed with iron deficiency (ID) (Short and Domagalski, 2013). ID is very common in children and women of LMIC, and occurs in ∼40% of otherwise healthy pregnant women in the US (Zimmermann, 2020) and up to 50% of healthy people who engage in endurance training (Epstein et al., 2018; Parks et al., 2017; Sinclair and Hinton, 2005). To explore the response to iron supplementation of the gut microbiota in healthy humans, we recruited 20 participants to provide daily stool samples for one week before, the one week during, and for one week after a 7-day period of daily ingestion of a commercial iron supplement (65 mg iron/325 mg ferrous sulfate). One participant (#8) had been taking iron supplements for many years, and hence sampled daily for 21 days with their normal iron supplementation to determine baseline fluctuations. Of the sixteen participants that completed the study, all provided at least 18 samples, resulting in 301 samples total (Figure 1A).

**Figure 1:**
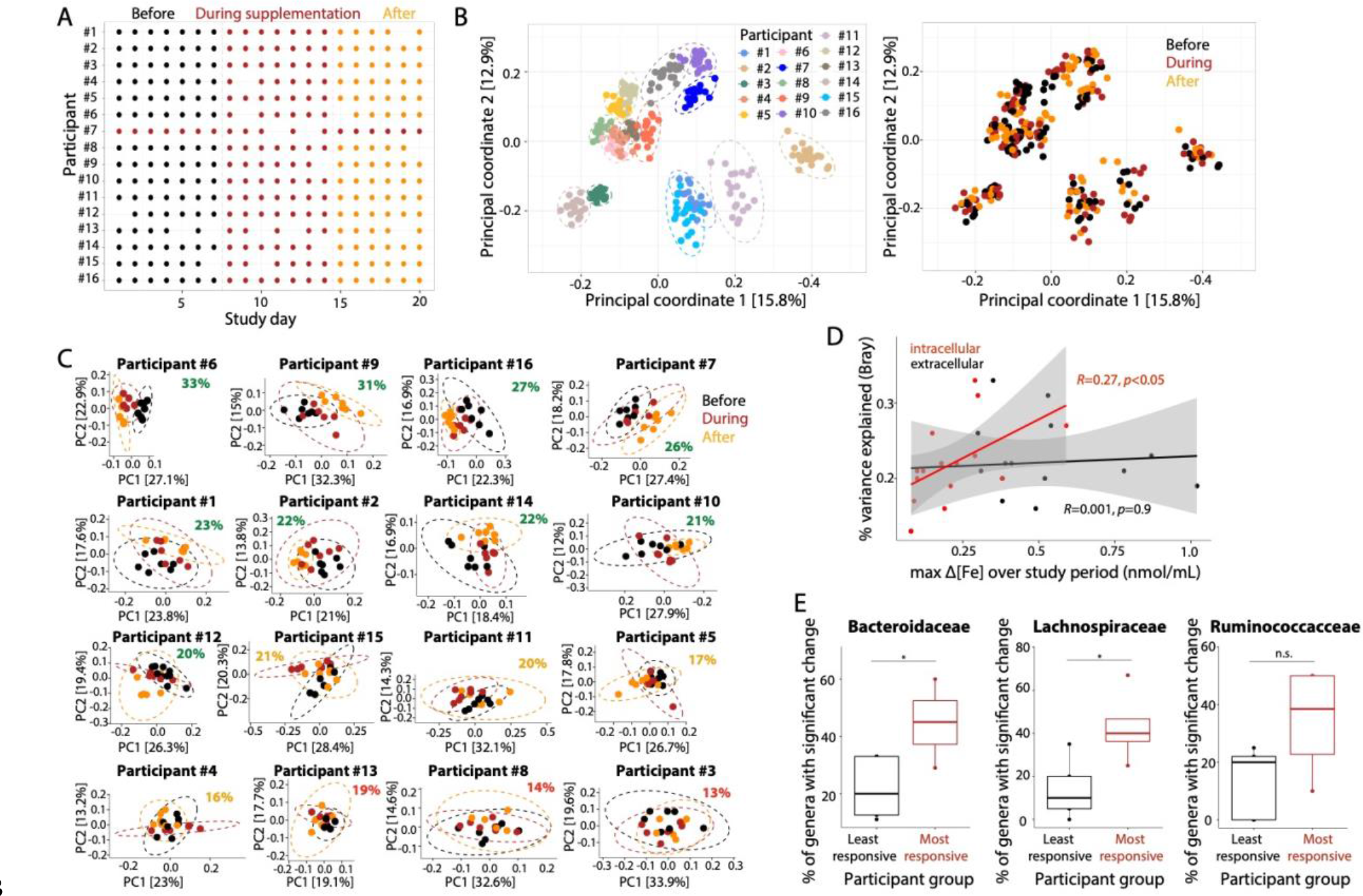
Iron supplementation results in participant-specific shifts in gut microbiota composition that are correlated with intracellular iron concentration. A) A total of 301 stool samples were collected from 16 participants asked to sample daily for 7 days before, 7 days during, and 7 days after a period of daily iron supplementation. All participants provided ≥18 samples. Fifteen participants supplemented with iron only for the 7 days; participant #8, a long-time user of iron supplements, took iron supplements all 21 days and thus served as a control for determining baseline fluctuations. B) The effects of iron supplementation on gut microbiota composition do not overcome inter-individual variation. A Principal Coordinate Analysis of pairwise Bray-Curtis distances clustered samples largely by participant as compared to time relative to iron supplementation. C) Examination of each participant individually showed that iron supplementation significantly accounted for 20-33% of the variability among samples in 9 participants (*p*<0.001, percent variability in green) and 16-21% in 4 participants (*p*<0.05, percent variability in yellow). Iron did not induce significant changes in the microbiota composition of participants #3 and #13 (*p*>0.05, percent variability in red). D) Intracellular iron concentrations measured in stool samples were correlated with percent variability in community structure (*R*=0.27, *p*<0.05), while extracellular concentrations showed no correlation (*R*=0.001, *p*=0.9). E) The percent of genera or species in the Lachnospiraceae and Bacteroidaceae families that exhibited significant changes during and after iron supplementation was significantly greater in the four participants that were most responsive to iron as compared with the participants that were least responsive to iron and control participant #8 (*: *p*<0.05, HSD test). n.s.: not significant.

We extracted DNA from each sample for 16S rRNA gene sequencing. Variation in community structure across all samples was visualized via Principal Coordinate Analysis (PCoA) of pairwise Bray-Curtis distances; other distance metrics led to similar conclusions (Figure S1). Samples largely clustered by participant (Figure 1B), indicating that the effect of iron supplementation on the gut microbiota at clinically relevant doses does not overcome to inter-individual variation. Indeed, iron supplementation explained only 2.8% of the variability in community structure throughout the study, while participant identity explained 87% (*p*<0.001, ANOVA). Nonetheless, when examining each participant of our current study individually, iron supplementation accounted for a much larger proportion of the variance: 20-33% for 9 participants (*p*< 0.001, ANOVA) and 16-21% for 4 of the remaining participants (*p*<0.05, ANOVA) (Figure 1C). For two participants (#3 and #13), iron did not induce significant changes in community structure. Thus, a normal dose of iron supplementation elicits small but detectable individual-specific changes in the gut microbiota of most healthy humans.

Given the high degree of participant specificity in the responses, we focused on subsets of the individuals to determine whether there were general structural changes to the gut microbiota. Among the 4 participants (#6, #7, #9, and #16) for which supplementation accounted for the largest proportions of variance, in participant #16, the gut community shifted during iron supplementation along one vector and continued to shift along that same vector after supplementation was halted (Figure 1C). Similar behavior was seen in participants #6, #7, and #9, suggesting that iron supplementation causes microbiota alterations that are either not reversible in the time frame of 1 week, or represent a delayed response to iron supplementation. In fact, iron-induced changes in community structure were of larger magnitude when comparing the “before” versus “after” periods (9 participants with *p*<0.05, 18-34% of total variance) than they were when comparing the “before” versus “during“ periods (4 participants with *p*< 0.05, 14-26%), again indicating that the effects of iron supplementation are gradual and delayed.

We wondered whether the participant-specific responses to iron supplementation were associated with the timing of iron distribution or the local concentrations of iron throughout various parts of the gastrointestinal tract. Bowel movement frequency was uncorrelated with the percent variability attributable to iron across participants (Figure S2), suggesting that transit time of the iron through the gut was not a major determinant of participant-specific changes. We quantified extracellular and intracellular iron concentrations of each sample by separating the supernatant from the bacterial pellets. While the magnitude of change in extracellular concentration was uncorrelated with percent variability (*R*=0.001, *p*= 0.9, F-test), intracellular iron concentration was correlated with differences in percent variability (*R*=0.27, *p*< 0.05, F-test) (Figure 1D), suggesting that participant-specific responses might be explained by the capability of members of the community to internalize exogenous iron.

### Participant-specific Lachnospiraceae and Bacteroidaceae genera are the most responsive to iron supplementation

To determine the taxonomic families most responsive to iron supplementation, we compared the relative abundance of several major families among the three study time periods (before, during, and after iron supplementation) in the four participants that were most responsive to iron supplementation (#6, #7, #9, and #16). The Akkermansiaceae, Enterococcaceae, and Enterobacteriaceae were present at very low abundances or not detected in most participants; hence, we focused on the Bacteroidaceae, Lachnospiracace, and Ruminococcaceae families. Although there were not consistent changes across the four participants at this taxonomic level (Figure S3), the relative abundance of certain genera or species affiliated with these families exhibited significant changes upon iron supplementation (*p*<0.05, HSD test), suggesting that iron-induced changes are most prominent at these lower taxonomic levels.

The week-to-week changes in our control participant that took supplements throughout the three weeks (#8) were small (14% of variation), and not statistically significant (*p*=0.169, PERMANOVA). Therefore, to identify structural changes that were specifically induced by iron supplementation, we calculated the fraction of genera/species that changed significantly within each family for each participant, and compared these fractions among the four participants that were most responsive to iron (26-33% variance explained, *p*<0.001) and among those that were least responsive to iron (#3,# 4, #13, and #5; 13-19% variance, *p*>0.05), as well as the control participant (#8). The percent of varying genera in the Bacteroidaceae and Lachnospiraceae families was significantly greater in the four participants most responsive to iron supplementation (Figure 1E), suggesting that genera/species within these two families in particular responded to iron supplementation and merit deeper understanding of their responses to iron levels.

### In vitro passaging and study of an iron-responsive community recapitulates the observation that iron supplementation induces only small changes in community composition

Given the inherent challenges in interpreting microbiota responses in the context of a complex host, we next sought to quantify the extent to which communities of gut commensals are sensitive to iron levels in a more controlled environment. Motivated by our previous study demonstrating that stool-derived *in vitro* communities can mimic the composition of the fecal microbiome and its sensitivity to ciprofloxacin observed *in vivo* (Aranda-Díaz *et al*., 2020), we used a pre-supplementation fecal sample from participant #16 (based on their relatively large response to iron) to derive *in vitro* communities via repeated passaging in BHIS+0.05% mucin (Figure 2A). After five 48-h passage cycles, the community was stable and contained ∼30 ASVs (Callahan et al., 2016), which collectively represented 86% of the families detected in the fecal sample including members of the Bacteroidaeceae, Ruminococcaceae, Lachnospiraceae, Enterobacteriaceae, and Enterococcaceae families, all of which were predicted to respond to iron from our *in vivo* study (Figure 2B). Thus, this community provides a powerful starting point for interrogating the response to iron levels *in vitro*.

**Figure 2:**
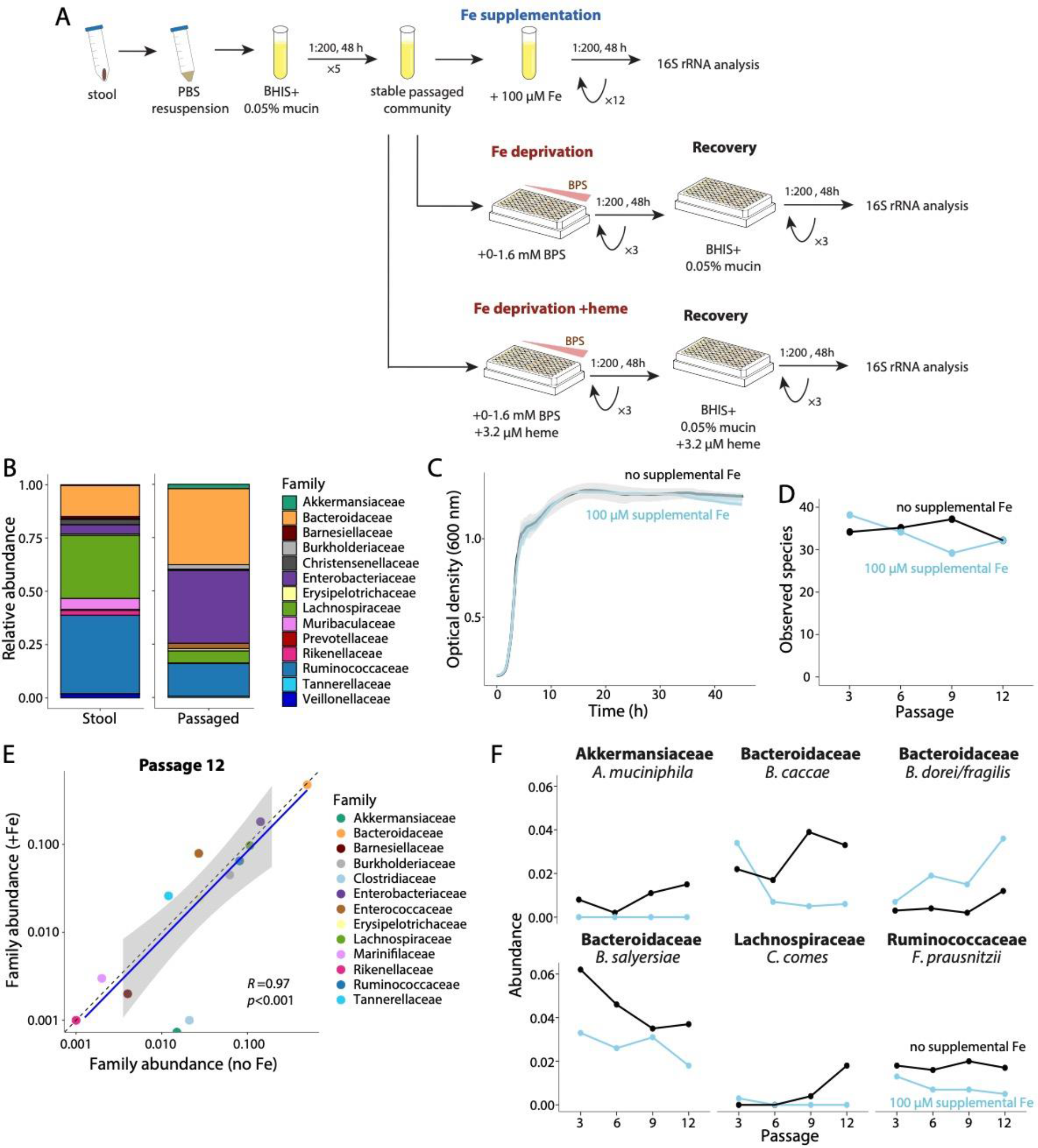
*In vitro* passaging of stool from an iron-responsive participant preserves microbial diversity and recapitulates the small changes in community composition due to iron supplementation observed *in vivo*. A) After 5 48-h passages, a community derived from a participant #16 pre- supplementation stool sample was passaged in BHIS+0.05% mucin medium either 12 times supplemented with 100 µM iron sulfate to study the effects of iron supplementation or 3 times supplemented with 0-1.6 mM BPS to study the effects of iron deprivation. Passaging was performed with and without heme to determine the degree to which heme can be used as an iron substitute. Iron- deprived communities were subsequently passaged 3 more times in the absence of BPS to assess recovery of the affected species. B) A stool-derived community from participant #16 retained 86% of the families (representing 89% of the relative abundance) detected in the stool sample after 5 48-h stabilization passage cycles. C) Addition of supplemental 100 µM iron sulfate did not affect the growth rate or carrying capacity of the *in vitro* community. Growth curves were measured after twelve 48-h passages in BHIS+0.05% mucin medium (without added heme). D) The number of observed species in the community supplemented with 100 µM iron sulfate was stable over passages and similar to that of the non- supplemented community. E) The relative abundance of most bacterial families after 12 passages was unchanged by iron supplementation (*R=*0.97), indicating that iron did not have large effects on community composition. F) Iron supplementation did not affect the relative abundance of most genera and species, except for small decreases in the relative abundance of *Faecalibacterium prausnitzii* (Ruminococcaceae), *Akkermansia muciniphila* (Akkermansiaceae), *Bacteroides salyersiae* (Bacteroidaceae), *Coprococcus_3 comes* (Lachnospiraceae) and *Bacteroides caccae* (Bacteroidaceae), and an increase in the relative abundance of *Bacteroides dorei/fragilis* (Bacteroidaceae).

To study the effects of iron supplementation on this community, we passaged the stabilized community 12 times in 96-well plates in BHIS+0.05% mucin medium (without added heme), with and without supplemental 100 µM iron sulfate (Figure 2A). The maximum growth rate and carrying capacity of the community were unaffected by supplemental iron (Figure 2C), indicating that a surplus of iron does not enhance the growth of at least the fastest growing species. To determine whether supplemental iron induced changes in community structure, we performed 16S rRNA gene sequencing on passages 3, 6, 9, and 12. The number of observed species remained largely stable over time and the number was similar in the supplemented and non-supplemented communities (Figure 2D), suggesting that the iron surplus did not have deleterious effects on most species. To determine the effects of iron supplementation on specific taxa, we examined the relative abundance of each major family over the 12 passages.

Consistent with observations from our clinical study, the supplemented and non-supplemented communities had very similar structure (Figure 2E, *R=*0.97), indicating that the community was insensitive to a surplus of iron at the family level.

Although the relative abundances of most genera/species were also similar between the supplemented and non-supplemented communities, a few species displayed substantial differences: *Bacteroides caccae* and *Bacteroides salyersiae* (Bacteroidaceae), *Faecalibacterium prausnitzii* (Ruminococcaceae), and *Coprococcus_3 comes* (Lachnospiraceae) were at lower abundances and *Akkermansia muciniphila* (Akkermansiaceae) was undetected in the iron-supplemented community compared to the non-supplemented community by passage 12 (Figure 2F). Interestingly, *Bacteroides dorei/fragilis* (Bacteroidaceae) was the only species for which supplemental iron had a positive effect (Figure 2F). Together, these results suggest that supplemental iron has small but measurable effects on community structure, which are isolated to a small group of species in the Bacteroidaceae, Ruminococcaceae, and Lachnospiraceae families. The similarities between our *in vitro* data and the findings of our *in vivo* study suggest that our *in vitro* community can be used more broadly to study the response to changes in iron levels.

### Lack of iron inhibits the growth of complex communities in vitro and irreversibly reduces community richness

To further probe how changes in iron concentrations affect human gut communities and to identify taxa that are most sensitive to iron availability, we next sought to characterize the effects of iron deprivation on our *in vitro* community. We passaged the stabilized community multiple times in 96-well plates in BHIS+0.05% mucin medium without added heme and supplemented with the extracellular iron chelator bathophenanthroline disulfonate (BPS) at increasing concentrations, thereby reducing iron levels in a graded fashion (Figure 2A). The maximum growth rate and carrying capacity of the community decreased in a chelator concentration-dependent fashion within a single 48-h passage (Figure 3A). Both growth parameters started to decrease substantially at 0.05 mM BPS, which is approximately the concentration necessary to chelate the 16 µM of iron in the medium given the 3:1 stoichiometry of BPS:Fe binding. The highest concentration of chelator (1.6 mM) reduced maximum growth rate and carrying capacity by 64% and 42%, respectively, suggesting severe inhibition of fast-growing species and potential extinction of some species (Figure 3A). Maintaining low iron conditions with BPS for two additional 48-h passages further reduced maximum growth rate and carrying capacity, with a total decrease of up to 88% and 67%, respectively, relative to the untreated community (Figure 3B,C), indicating gradual buildup of the effects of iron deprivation.

**Figure 3:**
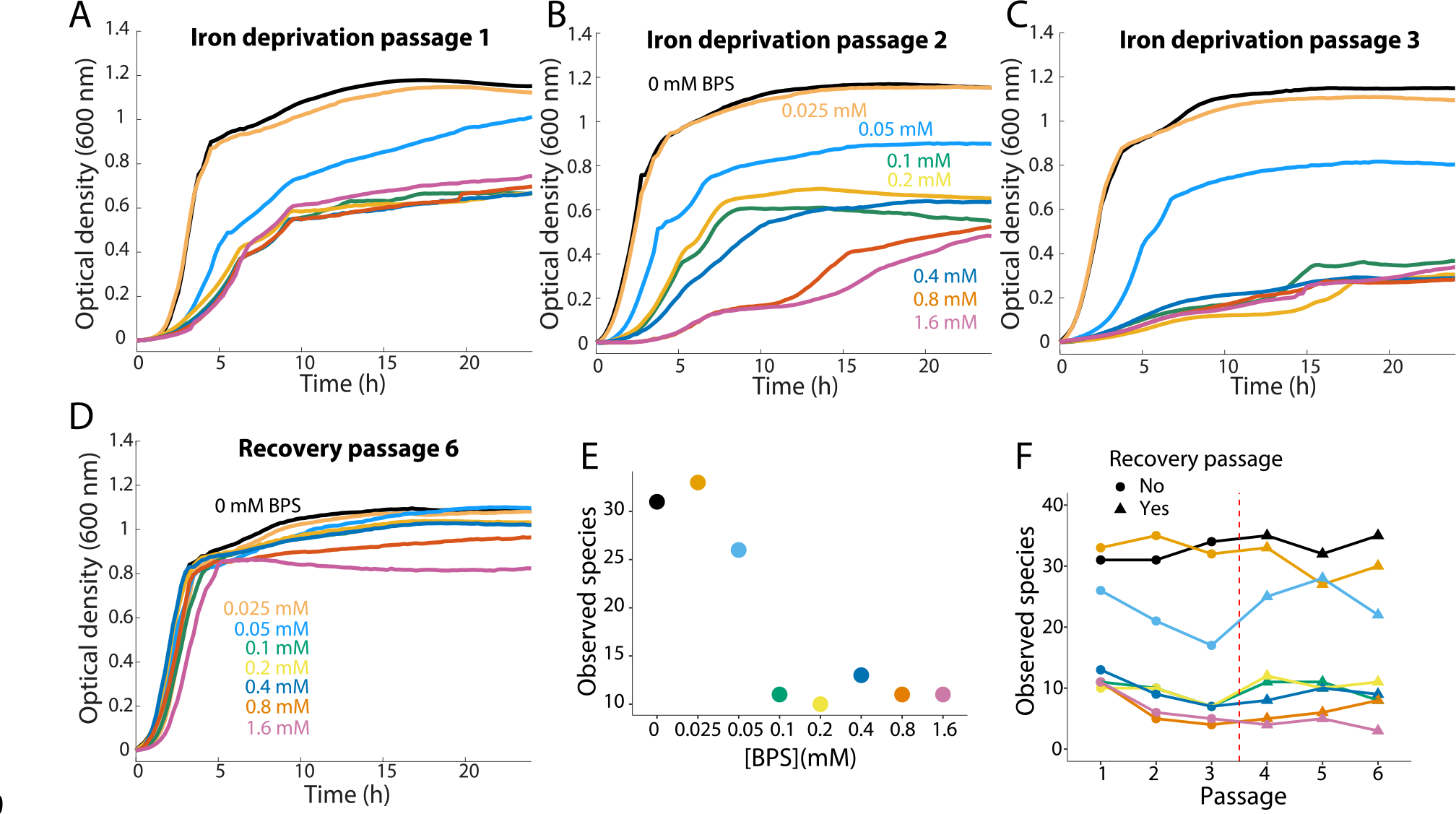
Iron deprivation decreases community yield and diversity. A) Growth rate and carrying capacity of the stool-derived community decreased in a BPS concentration-dependent manner in a single passage. B,C) The effects of iron deprivation were cumulative, as maintaining the communities in low-iron conditions further reduced growth rate and carrying capacity. D) The effects of iron deprivation were long lasting, as reintroducing the community to iron-sufficient levels restored maximum growth rate but not carrying capacity, suggesting permanent loss of species. E) The number of observed species decreased by ∼20-60% in a BPS concentration- dependent manner within a single 48-h passage. F) Maintenance of the community in iron-deprived media for 2 additional passages (passages 2, 3) further decreased the number of observed species, and the number of species did not recover after the community was reintroduced to iron- sufficient conditions (passages 4-6).

Reintroduction of the community to iron-sufficient concentrations (unaltered BHIS) restored maximum growth rate by the third 48-h recovery passage, suggesting that growth of the fastest-growing species fully recovered (Figure 3D). However, the carrying capacity of communities previously treated with BPS remained up to ∼30% lower than the untreated community, indicating either permanent loss of species whose niche remained unfilled or a change in metabolism that was irreversible within this time frame. Taken together, iron deficiency cumulatively destabilized human-derived *in vitro* communities over time and had long-lasting effects.

Motivated by the large, long-term changes in *in vitro* community growth due to iron deprivation, we performed 16S rRNA gene sequencing on all passages. The number of observed species in the community decreased in a BPS concentration-dependent fashion (Figure 3E), suggesting that the decrease in carrying capacity was at least partially due to severe depletion of some species. Reducing extracellular iron concentrations to nearly zero with 0.05 mM BPS decreased the number of observed species by ∼20% over the first 48-h passage and by ∼50% after a third 48-h passage relative to the untreated community (Figure 3F), suggesting that a large fraction of community members depend on iron to survive and/or grow and further emphasizing that the effects of iron deprivation are gradual and cumulative.

Virtually complete elimination of extracellular iron by adding excess chelator (BPS >0.1 mM) decreased the number of observed species to an even greater extent, by ∼40% over one 48-h passage and ∼80% by the third passage (Figure 3F), indicating variability in sensitivity and/or adaptation to low-iron conditions across species, whereby some could survive only at lower BPS concentrations. When the communities were reintroduced to iron-sufficient conditions, the number of observed species remained at approximately the same reduced level even after three 48-h recovery passages, suggesting irreversible loss of iron-dependent species (Figure 3F, passages 4-6).

### Sensitivity to iron deprivation is heterogeneous across taxonomic groups

We next assessed changes in community structure to identify the taxa that were strongly affected by iron deprivation. The most striking changes were observed in the Lachnospiracae and Ruminococcaceae families, whose relative abundances each decreased ∼10-fold (from ∼10% to ∼1%) with 0.05 mM BPS and decreased below the limit of detection with >0.1 mM BPS (Figure 4A, passages 1-3). When iron levels were restored, both families re-equilibrated at ∼50-60% of their initial abundances (∼5-6%) after treatment with 0.05 mM BPS, and remained undetectable in communities treated with >0.1 mM BPS (Figure 4A, passages 4-6). These dramatic changes suggest that certain members of these families are highly dependent on exogenous iron and lack sufficiently effective mechanisms of adaptation to low-iron conditions. Indeed, the genera within these families exhibited different sensitivities (Figure 4B), suggesting that adaptation mechanisms and/or overall sensitivity to low iron may be genus-specific.

**Figure 4:**
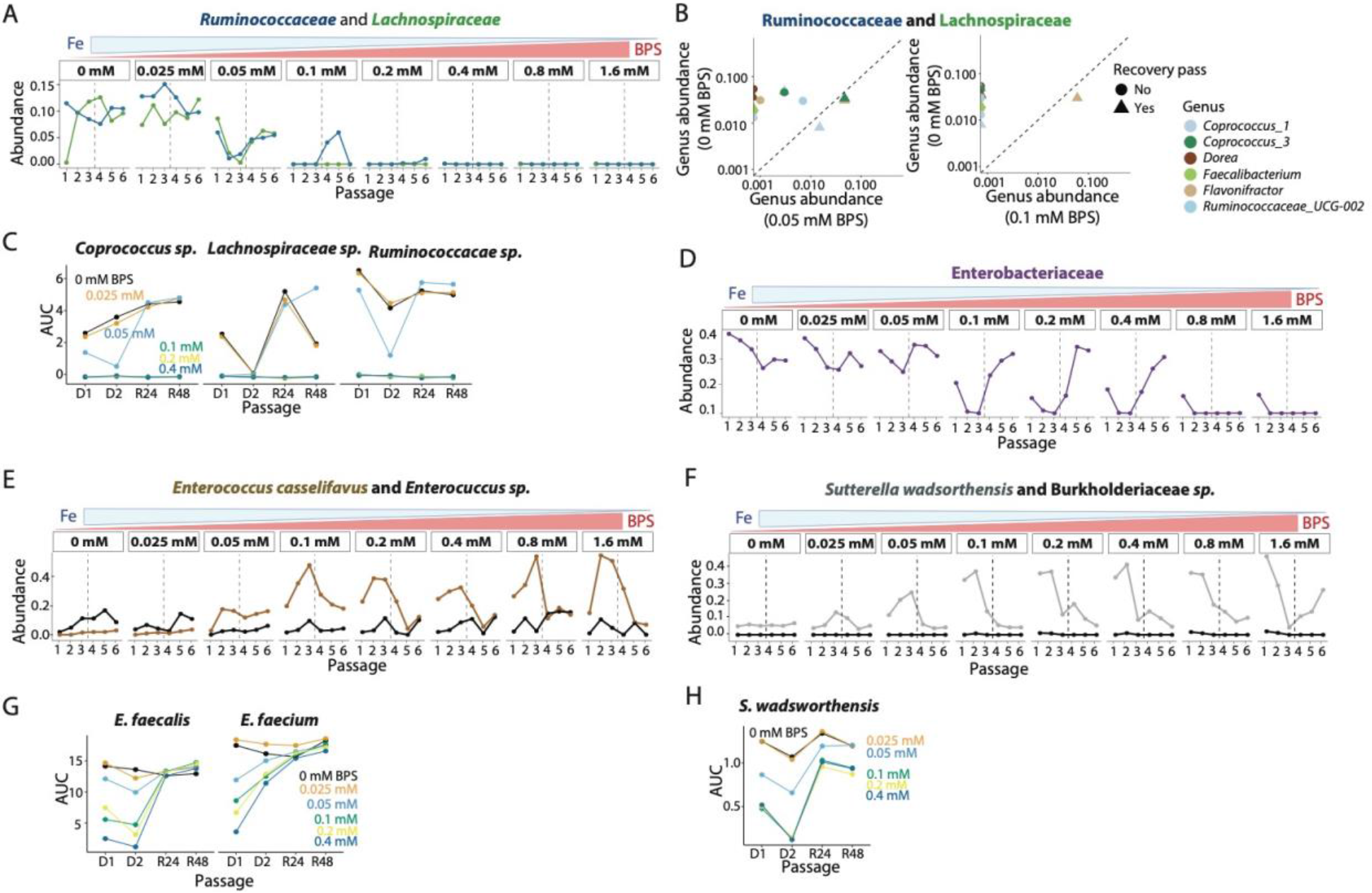
Sensitivity to iron deprivation is heterogeneous across families and across genera within a family. A) The Lachnospiraceae and Ruminococcaceae families were highly sensitive to iron deprivation. Both decreased below the limit of detection at >0.1 mM BPS during passages 1-3 and did not recover when iron-sufficient conditions were restored (passages 4-6). Vertical dotted line indicates the transition from iron-deprived to iron-sufficient conditions. B) *Flavonifractor* and *Coprococcus* species (members of the Lachnospiraceae and Ruminococcaceae families, respectively) were more resilient to iron deprivation (0.05 mM on left, 0.1 mM on right) than *Faecalibacterium* (Ruminococcaceae), *UCG-002* (Ruminococcaceae), and *Dorea* (Lachnospiraceae) species. C) A *Coprococcus* (Lachnospiraceae) species and other Lachnospiraceae and Ruminococcaceae isolates were highly sensitive to iron deprivation in isolated monocultures. Area under the curve (AUC) was calculated for each growth passage: iron-deprivation passage 1 and 2 (D1 and D2), recovery at 24 h of the second passage (R24), or recovery at 48 h of the second passage (R48). D) The Enterobacteriaceae family was sensitive to iron deprivation, decreasing below the limit of detection with >0.1 mM BPS. Nonetheless, Enterobacteriaceae relative abundance recovered in communities treated with ≤0.4 mM BPS when iron-sufficient conditions were restored. E, F) Enterococcaceae and Burkholderiaceae species increased in relative abundance during iron deprivation. G, H) In isolated monoculture, *E. faecium* (Enterococcaceae) was less sensitive to iron deprivation than *E. faecalis* (Enterococcaceae) and grew better in the second iron- deprivation passage than in the first passage. An *S. wadsworthensis* (Burkholderiaceae) isolate exhibited BPS concentration-dependent growth defects, but recovered effectively once iron-sufficient conditions were restored.

To determine whether sensitivity to iron deprivation was intrinsic to these families rather than an emergent property dependent on community context, we acquired human isolates of several Lachnospiraceae and Ruminococcaceae species and grew them in iron-deprived conditions (0-0.4 mM BPS) for two 48-h passages (Figure S5A). Consistent with their behavior in the community, the isolates exhibited species-specific growth defects at 0.05 mM BPS, and failed to grow after just one passage with ≥0.1 mM BPS (Figure 4C, S5B). These results suggest that overall sensitivity to iron deprivation is intrinsic to the Lachnospiraceae and Ruminococcaceae families, with subtle differences at the species level, and that isolate responses to iron deprivation can inform community-level changes.

The next most sensitive family to iron deprivation in the community was the Enterobacteriaceae family, which was represented by a single ASV in the *Escherichia*/*Shigella* genus. The relative abundance of this species decreased during iron deprivation in a BPS concentration-dependent manner and dropped below the limit of detection by the third 48-h passage with >0.1 mM BPS concentrations (Figure 4D, passages 1-3), suggesting that this commensal *Escherichia* species is highly dependent on exogenous iron. However, unlike the Lachnospiraceae and Ruminococcaceae families, when the community was reintroduced to iron-sufficient conditions, the *Escherichia* species fully recovered to its pre-treatment relative abundance by the third 48-h recovery passage in communities treated with ≤0.4 mM BPS (Figure 4D, passages 4-6), suggesting that this species possesses efficient low-iron adaptation mechanisms that allow it to endure long periods of iron restriction and then flourish once iron-sufficient conditions are restored. Notably, it did not recover in communities treated with >0.4 mM BPS, suggesting more complete chelation of iron atoms at these higher concentrations and indicating that the iron-binding molecules (siderophores) produced by this species have a lower affinity for iron than BPS, unlike the high-affinity siderophores produced by pathogenic *E. coli* (*K*=10^-52^ M^-1^) (Carrano and Raymond, 1979). Taken together, these results suggest that commensal *Escherichia* species have a higher dependency on iron than pathogenic species from the same genus but are equipped with efficient low-iron adaptation mechanisms that are likely missing in members of the Lachnospiraceae and Ruminococcaceae families.

Iron deprivation did not have detrimental effects on the Enterococcaceae and Burkholderiaceae families, which are low-abundance yet prevalent members of the human gut microbiome and are largely understudied. Most knowledge about the Enterococcaceae focuses on pathogenesis, and specifically on infection by the opportunistic pathogen *Enterococcus faecalis* (Keogh et al., 2016; Krawczyk et al., 2021; Saillant et al., 2021). Literature on iron responses of the Burkholderiaceae family is almost nonexistent (Hiippala et al., 2016). When iron concentrations were reduced (0.05 mM BPS) or eliminated with excess chelator (>0.1 mM BPS), Enterococcaceae relative abundance gradually increased in each passage, by ∼6-fold (from ∼10% to ∼60%) by the end of the third 48-h passage (Figure 4E, passages 1-3), suggesting that this family has little to no dependency on exogenous iron for survival and/or rapid, efficient mechanisms of adaptation to low iron. When iron-sufficient conditions were restored, Enterococcaceae relative abundance returned to near pre-treatment levels (Figure 4E, passages 4-6). Of the two *Enterococcus* species detected in the community, the initially less abundant *E. casseliflavus/faecium* (present at <1% in the untreated community) increased in relative abundance during iron deprivation, while the initially dominant species was at the same or lower relative abundance (Figure 4E), suggesting that Enterococcaceae members have different levels of resilience to low iron. Indeed, during growth in isolation, *E. faecium* was less sensitive to iron deprivation than *E. faecalis* and exhibited more growth during the second passage in iron-deprived media than in the first (Figure 4G and S5C). Both species immediately recovered when reintroduced to iron-sufficient medium regardless of the concentration of previous BPS treatment, suggesting that members of the Enterococcaceae family are highly resilient to iron deprivation.

The Burkholderiaceae family, which is represented by a *Sutterella wadsworthensis* ASV in the *in vitro* community, increased in relative abundance from ∼5% in the untreated community to ∼35-40% during the first passage with BPS (Figure 4F, passage 1).

However, in subsequent passages with BPS, Burkholderiaceae relative abundance gradually declined (Figure 4F, passages 2 and 3), and upon reintroduction to iron- sufficient conditions its relative abundance returned to pre-treatment levels. This behavior suggests that *S. wadsworthensis* does not require high levels of iron for survival. When grown in isolation, *S. wadsworthensis* exhibited a BPS concentration- dependent decrease in growth rate and carrying capacity (Figure 4H, S5D). When reintroduced to iron-sufficient media, *S. wadsworthensis* recovered quickly, although not completely and in a BPS concentration-dependent manner (Figure 4H, S5D), suggesting a relatively high resilience to iron deprivation but lower than that of *Enterococcus* species.

### Iron deprivation restructures the Bacteroides genus

Iron needs and acquisition strategies in commensal gut *Bacteroides* species have not been well characterized, but some species such as *B. thetaiotaomicron* and *B. vulgatus* are known to benefit from siderophores that are produced by pathogenic *E. coli* or by *Salmonella* species in low-iron conditions (Rocha and Krykunivsky, 2017; Zhu et al., 2020). How commensal *Bacteroides* species respond to low-iron conditions in the context of a complex community and in the absence of pathogenic bacteria has not been investigated. In our community, the Bacteroidaceae family overall behaved differently from all other families. Bacteroidaceae relative abundance was ∼30% during all three iron-deprivation passages at all BPS concentrations (Figure 5A, passages 1-3), suggesting that at least some family members have little dependency on iron or have efficient adaptation mechanisms that allow them to survive low-iron conditions. When the community was reintroduced to iron-sufficient conditions, Bacteroidaceae relative abundance increased in a BPS concentration-dependent manner (Figure 5A, passages 4- 6).

**Figure 5:**
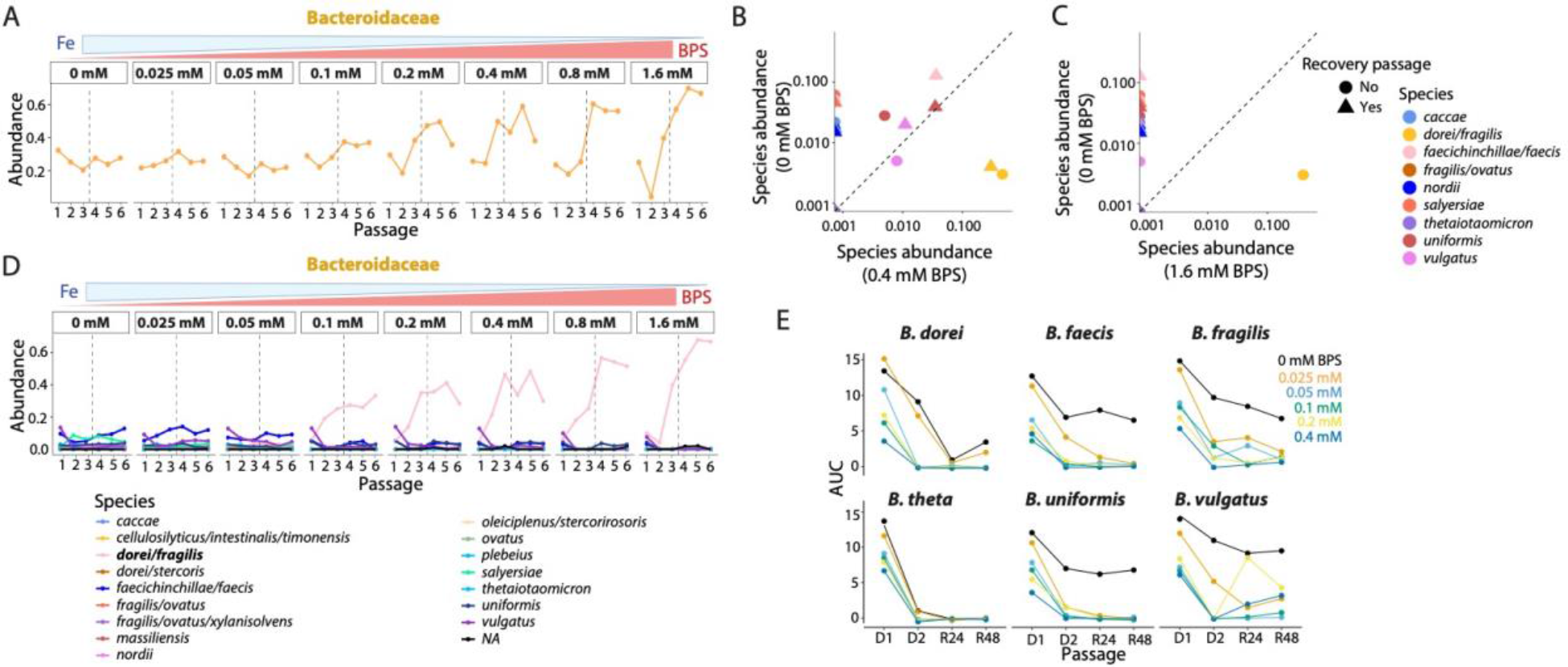
Iron deprivation can restructure community composition by decreasing competition. A) The Bacteroidaceae family maintained relatively constant relative abundance during iron deprivation, and increased in relative abundance during the recovery passages in a BPS concentration-dependent manner, suggesting that some member species that were less sensitive to iron expanded into niches left availably by more sensitive species. B,C) *Bacteroides* species exhibited distinct sensitivities to low iron (0.4 mM BPS in (B) and 1.6 mM in (C)). D) A *B. dorei/fragilis* ASV increased in relative abundance from <1% to ∼40% during iron deprivation and further increased to ∼65% during the subsequent iron recovery passages. E) *Bacteroides* isolates exhibited species-specific, BPS-concentration-dependent growth defects during iron deprivation. All species grew substantially more poorly in the second iron-deprivation passage even in the absence of BPS, suggesting that the absence of heme from our base medium inhibits growth after depletion of internal heme stores even in the presence of inorganic iron.

Further examination revealed that *Bacteroides* species have distinct sensitivities to iron deprivation. *B. faecichincillae/faecis*, *B. vulgatus*, and *B. uniformis* decreased in relative abundance from ∼5-10% to 0-1% during iron deprivation (Figure 5B), but recovered fully (*B. uniformis*) or partially (*B. faecichincillae/faecis* and *B. vulgatus*) when sufficient iron was restored (Figure 5B), suggesting that these species are iron-dependent but have adaptation mechanisms that allow them to survive. Importantly, this resilience disappeared for *B. uniformis* when communities were treated with >0.8 mM BPS or for *B. faecichincillae/faecis* and *B. vulgatus* when the communities were treated with >0.4 mM BPS, highlighting the variability in response to low iron within this family (Figure 5C).

Since 0.4 mM BPS is the concentration at which the *Escherichia* species became undetectable, it is also possible that these *Bacteroides* species depend on the production of siderophores by *Escherichia* or another species with similar behavior to meet their iron needs and/or a cross-feeding metabolite (Huus et al., 2021). *B. caccae, B. thetaiotamicron, B. nordii, B. salyersiae,* and *B. fragilis/ovatus*, which were all present at low (∼1%) but consistently detectable levels in the untreated community, became undetectable when iron concentrations were reduced and did not rebound once iron was restored to sufficient levels (Figure 5B and 5C), suggesting that these species are strongly dependent on iron and have less efficient adaptation mechanisms to low iron than *B. faecichincillae/faecis*, *B. vulgatus*, and *B. uniformis*.

Intriguingly, a *B. dorei/fragilis* ASV that was present at <1% in the untreated community increased in relative abundance during iron deprivation in a BPS concentration-dependent manner (Figure 5D, passages 1-3), comprising ∼40% of the community by the third 48-h iron-deprivation passage at the highest BPS concentration and increasing to ∼65% when sufficient iron levels were restored (Figure 5D, passages 4-6). The dominance of *B. dorei/fragilis* is presumably a combination of its lack of dependency on iron and/or efficient adaptation, with its ability to occupy the metabolic niches made available by species that did not survive the imposed low-iron conditions. Together, these results illustrate the dramatic extent to which differential iron sensitivities can substantially restructure the hierarchy within a single family.

A broad range of sensitivity to iron deprivation was also observed during isolated growth of six *Bacteroides* species in an initial passage in iron-deprived media (Figure 5E, S5E). During a second iron-deprivation passage, all *Bacteroides* species grew substantially worse than in the first passage or not at all (Figure 5E), again illustrating the gradual and cumulative effects of iron deprivation, as seen in the community (Figure 5B). However, the sensitivity of each species was more extreme in isolation than in the community, and growth defects were even observed in the absence of BPS, suggesting the absence of something other than iron from our BHIS base medium, which lacks heme. The need for exogenous heme (heme auxotrophy) in some *Bacteroides* species (e.g., *B. fragilis* and *B. thetaiotamicron*), and the potential for its supply by the human host or other members of the gut microbiota is well known (Gruss *et al*., 2012; Huus *et al*., 2021; Rocha and Krykunivsky, 2017). Thus, it is likely that the differences between growth in isolation versus in the community were at least partially due to the absence of heme. Because heme can be a source of iron, sensitivity to iron deprivation versus the requirement for heme as a cofactor are not easily disentangled in heme-deficient media. The substantial growth in the first passage without added heme and the gradual decline induced by iron deprivation suggest that *Bacteroides* species harbor heme and iron stores that enabled their transient growth and survival. Together, these community and isolate findings demonstrate why the inhibitory effects of iron deprivation are cumulative, and highlight the importance of heme as well as iron availability.

### Exogenous heme ameliorates the community destabilization induced by the lack of inorganic iron

Exogenous heme can be used as a cofactor and is directly incorporated into heme enzymes by heme auxotrophs. Moreover, heme is enzymatically broken down to iron by heme degradases in pathogenic bacteria (Wilks and Ikeda-Saito, 2014). The enzymes responsible for heme degradation in commensal bacteria have not been identified, and the prevalence of heme degradation for meeting inorganic iron needs is not well known. To determine whether heme is used as an alternate source of inorganic iron by the commensal bacteria in our community during iron deprivation, we repeated the passaging experiment with the inclusion of the 3.2 μM of heme that is usually present in BHIS medium. Although maximum growth rate and carrying capacity still decreased in a BPS concentration-dependent manner in the first 48-h passage, the presence of heme reduced the extent of the decreases in maximum growth rate and carrying capacity to 59% and 40%, respectively, at the highest BPS concentration (Figure 6A), suggesting that the fastest-growing species were less growth-inhibited, fewer species were lost, and/or that heme sustained the growth of heme auxotrophs that could not survive in its absence and that are less sensitive to low inorganic iron levels.

**Figure 6:**
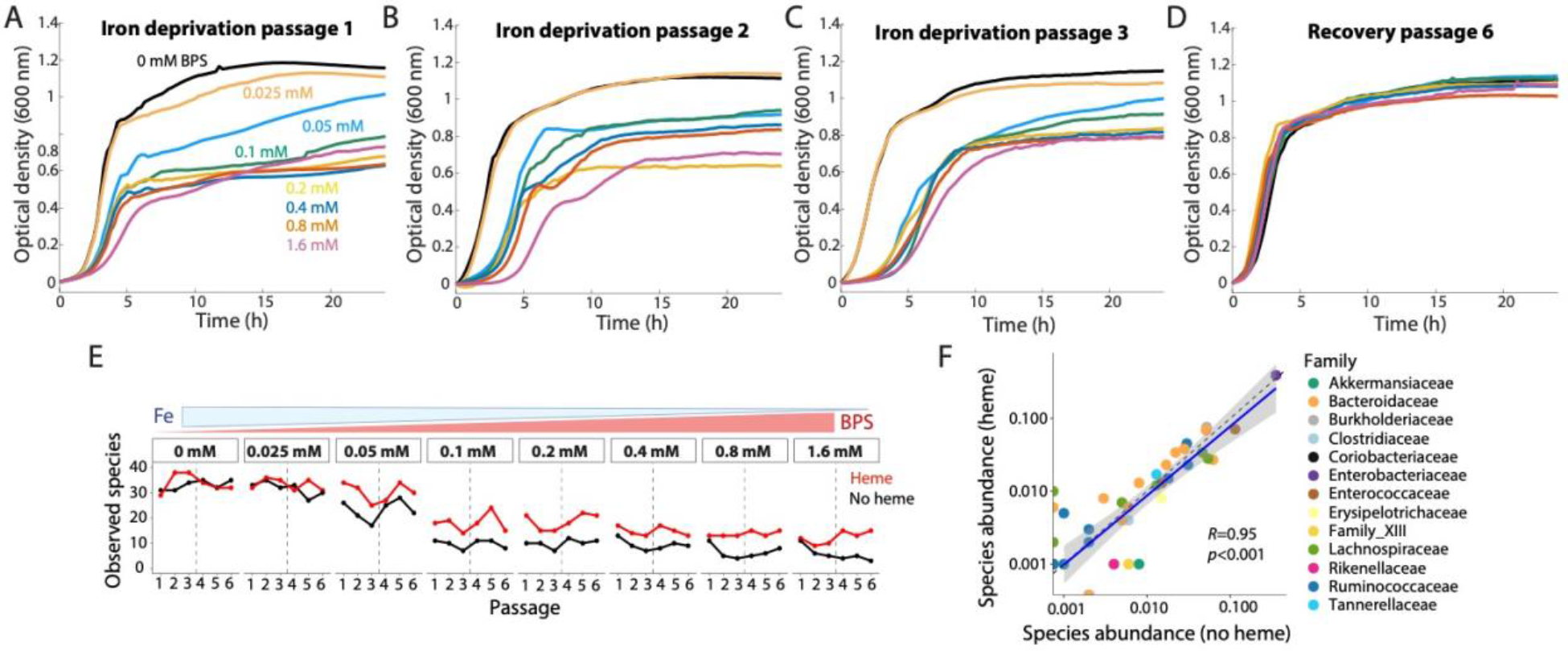
Heme supplementation buffers species extinction during iron deprivation. A-C) Supplementation with heme ameliorated the decreases in community growth rate and carrying capacity during iron deprivation, suggesting loss of fewer species and the use of heme as a source of iron. D) In the presence of heme, growth rate and carrying capacity were almost completely restored during the recovery passages. E, F) Addition of heme does not alter the number of observed species (E) or the relative abundance (F) of most species in the initial community (0 mM BPS) but prevents species extinction during iron deprivation (E, 0.05-1.6 mM BPS).

When maintaining the community in low-iron conditions for two additional passages, heme prevented further decreases in maximum growth rate and carrying capacity.

Instead, maximum growth rate was completely restored and carrying capacity was maintained by the third 48-h passage (Figure 6B and 6C), suggesting that heme was used as a source of iron by the fastest-growing species to restore their growth, and that the species that were sustained by heme in the first passage continued to thrive in low-iron conditions over an extended period. Reintroduction of iron to sufficient levels (unaltered BHIS) in the presence of heme restored growth rate and carrying capacity almost completely by the third 48-h passage (Figure 6D), suggesting that the depletion of species that are highly dependent on exogenous iron can be prevented by heme.

To determine whether the beneficial effects of heme during iron deprivation were due to its use as an alternate iron source or to its ability to sustain the growth of heme auxotrophs, we compared the number of observed species and the relative abundance of species in our community grown with and without heme. In iron-sufficient conditions, the number of observed species (Figure 6E, 0 mM BPS) and their relative abundance (Figure 6F) were very similar between the added-heme and no-heme communities, indicating that exogenous heme did not alter community structure in these conditions. However, when iron concentrations were reduced (0.05 mM BPS) or virtually eliminated (>0.1 mM BPS), exogenous heme reduced the number of species that were lost from ∼50% to ∼30% and from ∼80% to ∼60%, respectively, indicating that exogenous heme rescued some species that would otherwise have dropped below the limit of detection (Figure 6E). Together these results suggest that heme buffers the effects of iron deprivation by acting as an alternate source of inorganic iron.

### Heme rescues taxon depletion induced by iron deprivation in a family and species-specific manner

Given the buffering effects of heme during iron deprivation, we wanted to identify taxa that were rescued by heme and determine whether rescue was a family-level trait.

Heme was most advantageous to the *Escherichia* species in our community, almost completely preventing its drop in relative abundance induced by iron deprivation at ≤0.1 mM BPS and its apparent extinction up to the highest BPS concentration used (Figure 7A), suggesting that this commensal *Escherichia* species uses heme as an alternate source of iron and has efficient mechanisms for demetallating this cofactor.

**Figure 7:**
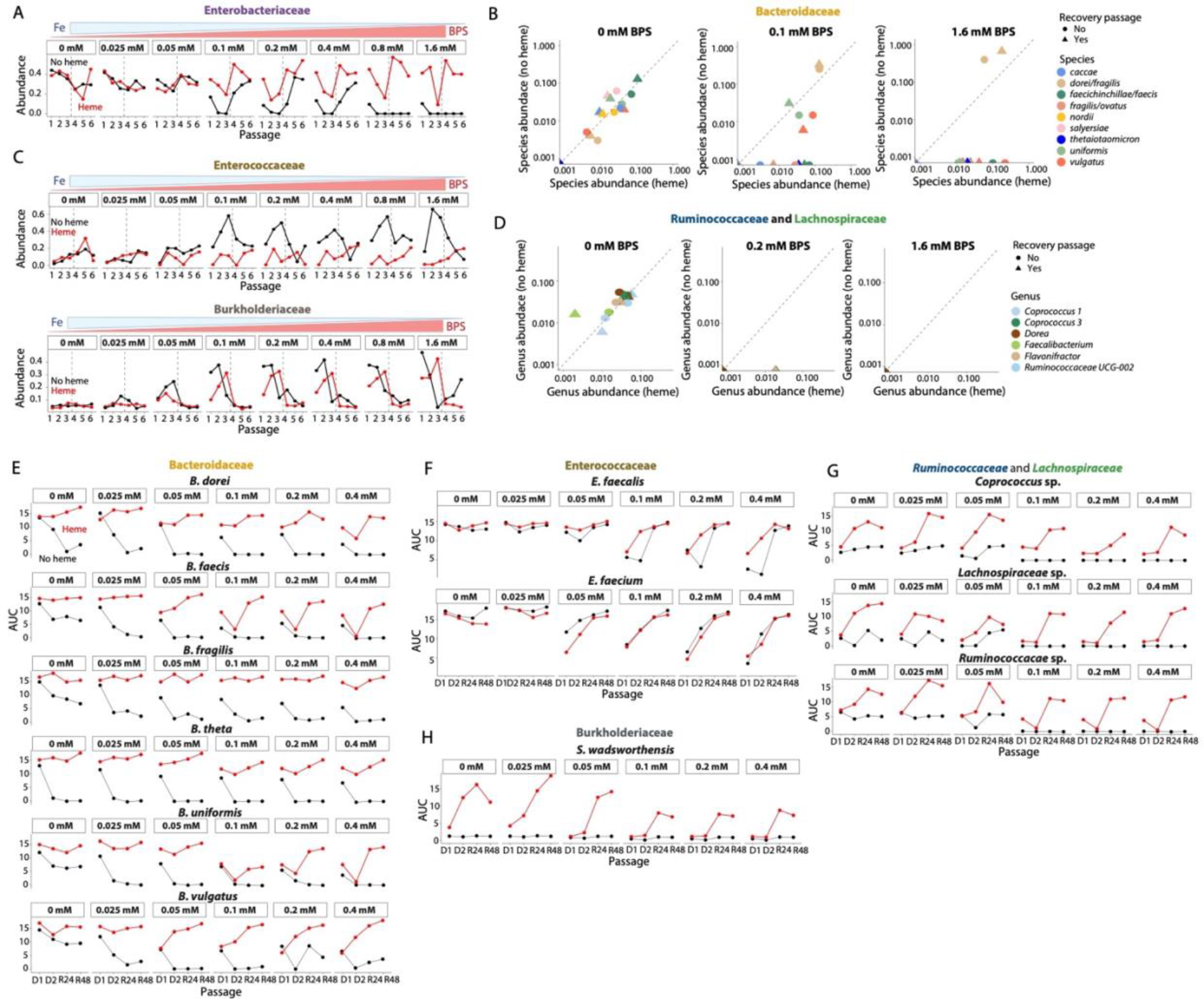
Rescue by heme during iron deprivation is species-specific. A) Heme ameliorated the decrease in relative abundance of the Enterobacteriaceae family at all BPS concentrations and even rescued it from extinction at 1.6 mM BPS, suggesting that its members have the ability to use heme as a source of iron. B) Heme completely prevents the extinction of all Bacteroidaceae species except *B. nordii* and *B. salyersiae* at 0.1 mM (middle) and 1.6 mM (right), indicating that the ability to use heme as an iron source is species-specific. Heme addition did not affect *Bacteroides* spp. relative abundances in the absence of BPS treatment (left). The *B. dorei/fragilis* ASV did not increase in relative abundance in the heme condition. C) The Enterococcaceae and Burkholderiaceae families did not increase in relative abundance in the presence of heme, suggesting that members of these families do not depend for heme-acquired iron for their survival. D) Of the genera in the Ruminococcaceae and Lachnospiraceae families, heme rescued only *Flavonifractor plautii* (Ruminococcaceae) and *Coprococcus_3 comes* (Lachnospiraceae) and only at ≤0.2 mM BPS (middle), suggesting that the ability to benefit from heme is species-specific and that species in these families do not use heme as efficiently as those in the Enterobacteriaceae (A) and Bacteroidaceae (B) families. Heme addition did not substantially affect the relative abundances of any genera in the absence of BPS treatment (left), and all species decreased below the limit of detection regardless of heme addition with 1.6 mM BPS (right). E) Heme ameliorated the negative effects of iron deprivation for all *Bacteroides* species during growth as isolated monocultures in a species-specific manner, with *B. fragilis* and *B. vulgatus* exhibiting the least growth defects in the presence of heme. The BPS concentration-dependent growth defects in the presence of heme suggest that these species need both inorganic iron and heme as a cofactor. F) For Enterococcaceae isolates, heme reduced the effects of iron deprivation for *E. faecalis* but had no effect on the growth of *E. faecium*. G,H) Heme rescued the iron deprivation-induced extinction of Lachnospiraceae, Ruminococcaceae, and Burkholderiaceae isolates and substantially enhanced their growth even in the absence of BPS, suggesting that heme may trigger alternative metabolic pathways in these species.

Heme also diminished the effects of iron deprivation and prevented the apparent extinction of all members of the Bacteroidaceae family except *B. nordii* and *B. salyersiae* (Figure 7B), suggesting that in addition to meeting the need of heme auxotrophs, heme can serve as an iron source for some but not all species in this family. Additionally, the relative abundance of *B. thetaiotaomicron*, *B. vulgatus*, and *B. fragilis/ovatus* actually increased when communities were iron-deprived in the presence of heme, indicating that heme provides an additional growth benefit to these species compared to inorganic iron. In accordance with our hypothesis that the *B. dorei/fragilis* species has little dependency on iron and occupies niches that become available, this species was unable to expand (Figure 7B) to the same extent as in the absence of heme (Figure 5B,C), consistent with the continued presence of members that became undetected in the non-heme community. Similar behavior was observed for the *Sutterella* (Burkholderiaceae) and Enterococcaceae species in this community (Figure 7C), suggesting that these families do not benefit from heme as an alternate source of iron and that their resilience to low iron depends on other adaptation mechanisms.

For the Lachnospiraceae and Ruminococcaceae families, heme enabled the survival at higher BPS concentrations of only one species in each family: *Flavonifractor plautii* (Ruminococcaceae) and *Coprococcus_3 comes* (Lachnospiraceae). However, heme did not prevent their eventual drop to undetectability (Figure 7D). These results suggest that heme can only act as an alternate source of iron for certain species within the Lachnospiraceae and Ruminococcaceae families, and their mechanisms for demetallating heme to obtain iron are not as efficient as those of the Bacteroidaceae and Enterobacteriaceae families.

Iron deprivation of isolates in the presence of heme largely resulted in similar behaviors as with communities. The growth rate and carrying capacity of all *Bacteroides* isolates decreased in a BPS concentration-dependent manner, and decreases were more pronounced in the second iron-deprivation passage, indicating the need for inorganic iron and not just heme as a cofactor (Figure 7E). However, heme ameliorated the negative effects of iron deprivation and enabled full or nearly full recovery when reintroduced to iron-sufficient conditions. Responses were species-specific: *B. fragilis* and *B. vulgatus* had the lowest growth defects and recovered fully when reintroduced to iron-sufficient conditions, while *B. thetaiotamicron*, *B. uniforms*, *B. faecis*, and *B. dorei* exhibited larger growth defects that lasted throughout the recovery passages. For Enterococcaceae species, heme reduced the effects of iron deprivation for *E. faecalis* but had no effect on the growth of *E. faecium* (Figure 7F).

In isolation, heme rescued Lachnospiraceae and Rumninococcaceae species from extinction due to iron deprivation at all BPS concentrations (Figure 7G). In addition, their growth was actually enhanced relative to growth in the absence of heme at all BPS concentrations, starting in the second iron-deprivation passage and lasting throughout the recovery passages (Figure 7G). Similar behavior was observed with *S. wadsworthensis* (Figure 7H). Although we currently cannot explain these positive effects on growth, it may be due to heme increasing expression of virulence genes and a shift from fermentation to more efficient anaerobic respiration, as has been reported for opportunistic pathogens (Palmer and Skaar, 2016; Yamamoto et al., 2005).

These findings highlight the varied, taxon-specific roles of heme on isolate and community growth during iron deprivation.

## Discussion

Here, we showed that a typical clinical dose of supplemental iron has gradual, long-lasting effects on the healthy human gut microbiota. The magnitudes of the effects were participant-specific. The low degree of variability due to iron supplementation (Figure 1B) was much smaller than the variability attributed to a five-day course of ciprofloxacin (antibiotics accounted for ∼14% of the variability and participant identity accounted for 67%, versus 2.8% and 87%, respectively, for iron supplementation) (unpublished work; Dethlefsen, Relman, et al.). The variation in responses across participants was partially due to differences in microbiota membership and the ability of these microbes to respond to increased iron via internalization (Figure 1D).

Nonetheless, across our cohort the most responsive taxa were members of the Bacteroidaceae and Lachnospiraceae families (Figure 1E), with the specific genera/species varying by participant. Of note, the relative abundance of the Enterobacteriaceae family, which had a prevalence of 66%, did not increase significantly with iron supplementation in any of our participants, suggesting that the *Escherichia* species in our healthy cohort are distinct from pathogenic *E. coli*, which is notable in its voracity for iron (Andrews *et al*., 2003; Galardini *et al*., 2020). *In vitro* communities derived from the stool of multiple participants in our study maintained most of the taxonomic diversity of the inocula (Figure 2B, S4), and iron supplementation of these *in vitro* communities yielded similar results as supplementation *in vivo* (Figure 2).

Moreover, these findings further illustrate the ability of *in vitro* communities to model the effects of environmental perturbations on the gut microbiota *in vivo*.

Stool-derived *in vitro* communities provided an opportunity to interrogate the global effects of iron levels on gut commensal communities, to gain a more nuanced understanding of the differences in iron sensitivities across taxa, and to identify commensals that benefit from heme as an alternate source of iron. Importantly, while we largely focused on a community derived from the participant that responded most strongly to iron supplementation, a community derived from another responder exhibited similar behaviors under changes to iron levels (Figure S4). By depriving *in vitro* communities of iron with the cell-impermeable chelator BPS, we showed that iron deprivation irreversibly reduces community richness, similar to previous studies in mice (Coe *et al*., 2021).

The environmental tunability of communities *in vitro* enabled us to discover that species depletion was BPS concentration-dependent and species-specific (Figure 3, 4, 5), indicating that commensal taxa have differential resilience to low-iron conditions, likely reflecting different iron needs and/or adaptation mechanisms. Members of the community covered the entire spectrum of sensitivity to resilience: Lachnospiraceae and Ruminococcaceae members were the most sensitive to low iron and rapidly became undetected (Figure 4A), Enterobacteriaceae were sensitive but quickly recovered when iron-sufficient conditions were restored (Figure 4D), *Sutterella* (Burkholderiaceae) and *Enterococcus* species (Enterococcaceae) were resilient to low iron (Figures 4E and 4F), and members of the *Bacteroides* genus spanned the spectrum in a species-specific manner (Figure 5). Moreover, heme buffered the effects of iron deprivation, preventing the extinction of species (Figure 6), and the ability to use heme as an alternate source of iron was species-specific (Figure 7). Different responses to varying iron concentrations within a genus created the potential for certain resilient species to become dominant when other species with overlapping niches were inhibited (Figures 4E, 5D), indicating that the diverse range of responses has the potential to restructure a community under conditions of iron limitation. Thus, future predictions of the response of a given gut microbiota will benefit from further characterization of the behavior of individual species, in both isolated and community contexts, particularly the mechanisms of adaptation to changing iron conditions. When iron is in excess, the expression of iron-storage proteins and antioxidant enzymes may be quickly and efficiently upregulated in some taxa, while it may be slow or completely absent in others (Abdul-Tehrani *et al*., 1999; Andrews, 1998; Andrews *et al*., 2003). To avoid deleterious effects of iron deprivation, some commensals may release iron-binding molecules that can be shared, while others shift expression of iron-dependent enzymes (e.g., ferredoxin or iron-superoxide dismutase) to enzymes that do not require iron (e.g., flavodoxin or manganese-superoxide dismutase) (Andrews, 1998; Andrews *et al*., 2003; Campbell et al., 2019; Kortman *et al*., 2014; Kramer et al., 2020).

In the future, mechanistic studies with bacterial isolates and mutant libraries (Liu et al., 2021; Price et al., 2018) may provide greater understanding of these responses and adaptation mechanisms, as well as provide the opportunity to engineer communities through manipulation of iron levels and sensitivities. Utilization of heme as both an iron source and a cofactor may be intertwined in low-iron conditions for some species, as they appear to be for *Bacteroides* species (Figure 4). Moreover, in some cases the effects of iron or heme deprivation are not immediately apparent but instead require multiple passages in low-iron conditions (Figure 4E), indicating that these species can store iron/heme and then utilize these stores when iron/heme is lacking. This storage may explain the gradual changes observed during iron supplementation *in vivo* (Figure 1C) and deprivation *in vitro* (Figure 3B-D). In some cases, species such as the *Bacteroides* were more sensitive in isolation than in the community (Figure 5E), reflecting the interaction of iron availability with sharing of other resources such as heme.

This study provides a foundation for further interrogation of community restructuring due to environmental perturbations. Our study suggests that even in the case of chemicals such as iron that affect the host and microbiota alike, *in vitro* bacterial communities can provide key insight into the microbiota response *in vivo*. For iron specifically, the taxon specificity of most responses suggests that metagenomic sequencing may prove useful for revealing whether changes to iron concentration selects for particular strains and substrains. Our findings motivate biochemical identification and characterization of the enzymes that anaerobically extract iron from heme in *Bacteroides* and other species; currently only one anaerobic heme degradase has been identified, in *E. coli* (LaMattina et al., 2017). Use of transposon mutant libraries (Liu *et al*., 2021; Wetmore et al., 2015) when iron conditions are perturbed could shed light on genotype-phenotype relationships, and the structural underpinnings of enzyme function could be facilitated by recent advances in deep learning-based protein folding (Jumper et al., 2021). Our approach combining *in vivo* and *in vitro* interrogation can also be applied to future human studies of supplementation of anemic participants and of supplementation with heme instead of ferrous sulfate, both of which should inform potential treatment for anemic patients. Finally, clinical investigations of whether fecal microbiota transplants can reverse the effects of iron deprivation on gut communities should be facilitated by the relative ease and throughput of testing responses of stool- derived *in vitro* communities, which can be used to tune conditions before translating to mice and humans.

## Methods

### Non-clinical iron supplementation study recruitment and sampling

Twenty healthy adults (39.8± 12.6 years of age) who had not taken antibiotics or been diagnosed with iron deficiency within the previous year were recruited to provide daily stool samples for 7 days before, 7 days during, and 7 days after daily intake of a 65 mg iron supplement (325 mg ferrous sulfate, Nature Made®). Sample collection consisted of ∼50 mg of stool being resuspended in 500 μL of DNA/RNA shield (Zymo Research, R1100) and ∼5 g being placed in empty sterile plastic tubes, separately. One participant had been taking iron supplements for years, and hence was sampled daily for 21 days without additional supplementation but continuing their regular supplementation to determine baseline fluctuations. Participants collected samples themselves and were instructed to immediately freeze the samples at home until the samples were brought to the laboratory for storage at -80 °C, which happened at the end of the sampling period. Fifteen out of 20 participants completed the study, with 14 participants providing at least 18 samples, resulting in 301 samples total (Figure 1A). Samples stored in DNA/RNA shield were used for 16S rRNA sequencing analysis. Aliquots of the 5 g samples were used for iron quantification and for *in vitro* community inoculation.

Informed consent was obtained from participants and the study protocol (#25268) was approved by the Administrative Panel for Medical Research on Human Subjects (Institutional Review Board) of Stanford University.

### DNA extraction and 16S rRNA library preparation

For human stool, samples stored in DNA/RNA shield were thawed on ice and vigorously mixed. Two hundred fifty microliters of each sample were transferred into 96-well PowerBead Plates (Qiagen, 27500-4-EP-BP). Seven hundred fifty microliters of RLT buffer from an AllPrep DNA/RNA 96 Kit (Qiagen, 80311) were added to each well and subsequent steps of DNA extraction were followed using manufacturer’s instructions. For *in vitro* communities, 50 µL of saturated bacterial cultures were extracted using a DNeasy UltraClean 96 Microbial Kit (Qiagen, 10196-4).

Three microliters of extracted DNA were used in 75-µL PCR reactions containing Earth Microbiome Project-recommended 515F/806R primer pairs (0.4 µM final concentration) and 5PRIME HotMasterMix (Quantabio, 2200410) to generate V4 region 16S rRNA amplicons. BSA at a final concentration of 100 ng/µL was also added to extracted DNA from stool samples. Thermocycler conditions were: 94 °C for 3 min, 35 cycles of [94 °C for 45 s, 50 °C for 60 s, and 72 °C for 90 s], then 72 °C for 10 min. PCR products were individually cleaned up and quantified using the UltraClean 96 PCR Cleanup Kit (Qiagen 12596-4) and the Quant-iT dsDNA High Sensitivity Assay kit (Invitrogen Q33120) before 200 ng of PCR product for each sample were manually pooled. Pooled libraries were sequenced with 250- or 300-bp paired-end reads on a MiSeq (Illumina).

### 16S rRNA data analysis

Samples were demultiplexed with QIIME2 v. 2021.2 (Caporaso et al., 2010) and subsequent processing was performed using DADA2 (Callahan *et al*., 2016). truncLeF and truncLenR parameters were set to 240 and 180, respectively, and the pooling option parameter was set to “pool=FALSE” unless otherwise indicated. All other parameters were set to the default. Taxonomies of the resulting ASVs were assigned using the assignTaxonomy function and the Silva reference database.

Downstream analyses were conducted in R studio. Data were largely manipulated using the phyloseq package. PERMANOVA analyses were performed using the vegan package and HSD-tests using the agricolae package.

### Iron quantification

A colorimetric ferrozine-based assay protocol previously used for quantification of iron in cultured astrocyte cells (Riemer et al., 2004) was modified and used to measure extracellular and intracellular iron concentrations in stool samples. In brief, ∼100 mg of stool were resuspended in 1 mL of TE buffer (10 mM Tris-HCl [pH 8], 1 mM EDTA) and vortexed vigorously. Resuspensions were centrifuged (6,000 rpm, 5 min) and 100 µL of the supernatant were transferred to a 2-mL 96-well plate for extracellular iron quantification. Pellets were washed twice with PBS, resuspended in 500 µL of lysis buffer (20 mM Tris-HC [pH 8], 2 mM EDTA, 1.2% TritonX-100, 20 mg/mL lysozyme), and incubated at 37 °C for 30 min. Cell debris was pelleted (13,000 rpm, 10 min) and 100 µL of cell lysate were transferred to a 2-mL 96-well plate for intracellular iron quantification. To release all protein-bound iron and cofactor-bound iron, 100 µL of 10 mM HCl and 100 µL of iron releasing agent (1:1 (v/v) 1.4 M HCl and 4.5 % (w/v) KMnO_4_) were added to each of the intra/extracellular samples and incubated at 60 °C for 2 h. After samples were cooled to room temperature, 30 µL of iron detection reagent (6.5 mM ferrozine, 6.5 mM neocuproine, 2.5 M ammonium acetate, 1 M ascorbic acid in dH_2_O) were added and mixtures sat for 30 min at room temperature. One hundred fifty microliters of this solution were transferred to a 96-well polystyrene microplate and absorbance was measured at 550 nm using an Epoch2 plate reader (BioTek). Iron concentrations were calculated by comparing sample absorbances to those of a range of FeCl_3_ concentrations of equal volume and prepared in a similar manner as the stool samples (mixtures of 100 µL of FeCl_3_ standard in 10 mM HCl, 100 µL NaOH, 100 µL iron releasing agent). Iron concentrations were normalized against stool sample wet weight.

### Human-derived in vitro community inoculation and passaging

We chose a pre-supplementation stool sample participants #16 and #7 from which to separately derive *in vitro* communities based on these participants exhibiting large responses to iron supplementation. Approximately 100 mg of stool was homogenized in 1 mL of PBS in an anaerobic chamber. After large particulates were allowed to settle (∼2 min), the mixture was diluted 1:100 in PBS, used to inoculate 3 mL of BHIS (BHI, 3.2 µM hemin (referred to as ‘heme’ in this study), 0.5 mg/mL L-cysteine, 5 µg/mL Vitamin K)+0.05% mucin medium (1:100), and incubated at 37 °C for 48 h. This culture was subsequently passaged (1:200) in test tubes every 48 h for a total of 5 passages to obtain a stable community composition.

To study the effects of iron supplementation, 1 µL of the stable community was used to inoculate 199 µL of BHIS+0.05% mucin medium containing no heme and 100 µM FeSO_4_ (100 mM stock freshly prepared in 10 mM HCl to prevent oxidation). Optical density at 600 nm (OD_600_) was measured in a 96-well polystyrene microplate using an Epoch2 plate reader (BioTek) with continuous shaking every 15 min. The community was passaged with 1:200 dilution in this iron-supplemented medium every 48 h for 12 passages.

To study the effects of deprivation, 1 µL of the stable community was used to inoculate 199 µL of BHIS+0.05% mucin medium, with or without heme, containing 0-1.6 mM bathophenanthroline disulfonic acid disodium salt hydrate (BPS, Sigma Cat. #146617) (Figure 2A). OD_600_ was measured as before and the community was passaged in BPS-containing media 3 times (treatment passages 1-3). After 3 iron-deprivation passages, the community was passaged 3 more times in BHIS+0.05% mucin medium with or without heme in the absence of chelator (recovery passages 4-6).

### Measurements of gut commensal isolate growth

Twelve human bacterial isolates from five taxonomic families were chosen for measurements of growth in axenic culture. Eight were from obtained from BEI Resources as part of their Human Microbiome Project collection: *Coprococcus* sp. HPP0048 (Lachnospiraceae), *Lachnospiraceae* sp. *1_4_56FAA* (Lachnospiraceae), *Ruminococcaceae* sp. *D16* (Ruminococcaceae), *Enterococcus faecium* TX1330 and *Enterococcus faecalis* TX1322 (Enterococcaceae), *Sutterella wadsworthensis* HGA0223 (Burkholderiaceae), and *Bacteroides thetaiotaomicron* CL09T03C10 and *Bacteroides dorei* CL02T00C15 (Bacteroidaceae). *Bacteroides faecis*, *Bacteroides fragilis*, *Bacteroides uniformis*, and *Bacteroides vulgatus* (Bacteroidaceae) were isolated from a stool sample from one of our study participants.

All isolates were streaked on BHIS+0.05% mucin agar plates and allowed to grow for 48-72 h at 37 °C in an anerobic chamber. For each strain, 3 mL of BHIS+0.05% mucin liquid medium were inoculated with a colony and grown for 48 h at 37°C. One microliter of saturated culture was used to inoculate 199 µL of BHIS+0.05% mucin medium, with or without heme, containing 0-0.4 mM BPS (Figure S5A). OD_600_ was measured as before. At the end of the first 48 h passage, cultures were passaged in this BPS-containing media for another 48 h. After 24 h and 48 h of the second passage, 1 µL was used to inoculate 199 µL of BHIS+0.05% mucin medium, with or without heme, without chelator and OD_600_ was measured thereafter to access recovery at these two growth time points.

## Author Contributions

A.I.C., D.R., and K.C.H. designed the research; A.I.C. performed the research; A.I.C. analyzed the data; and A.I.C., D.R., and K.C.H. wrote the paper. All authors reviewed the paper before submission.

## Acknowledgements

The authors are grateful to members of the Relman lab and Huang lab for helpful discussions, with a special thanks to Katherine Xue, Handuo Shi, and Les Dethlefsen for their thoughtful comments on the manuscript, and to all the participants who made our clinical study possible. The authors acknowledge funding from the Stanford Microbiome Therapies Initiative (to A.I.C., D.A.R., and K.C.H.), the Thomas C. and Joan M. Merigan Endowment at Stanford University (to D.A.R.), NIH RM1 Award GM135102 (to K.C.H.), and NIH R01 AI147023 (to D.A.R. and K.C.H.). D.A.R. and K.C.H. are Chan Zuckerberg BioHub Investigators.

## Supplementary Figures

**Figure S1:**
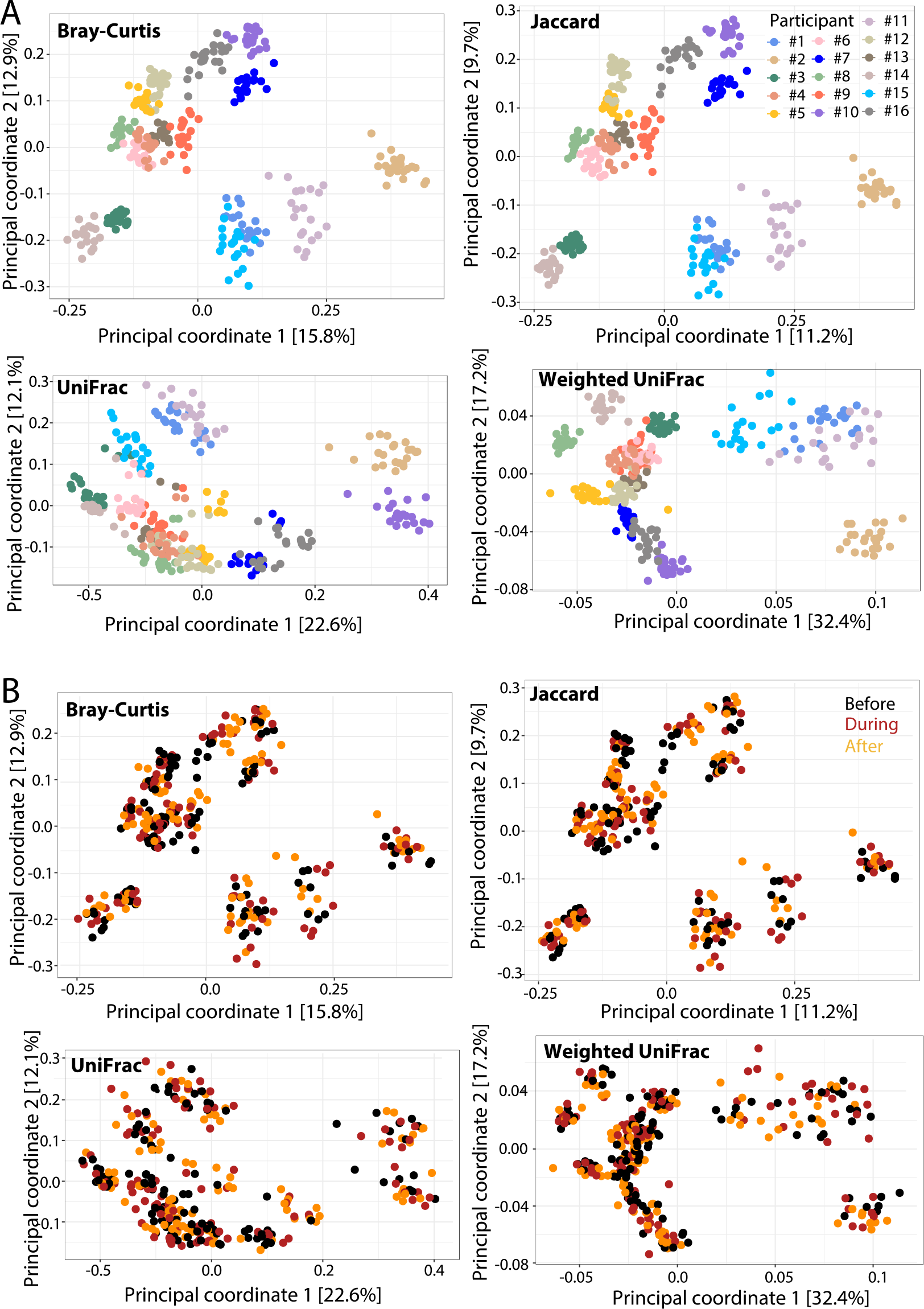
Multiple beta diversity metrics show that samples cluster largely by participant. A,B) Principal Coordinate Analyses (PCoA) of pairwise Bray-Curtis, Jaccard, unweighted UniFrac, or weighted UniFrac distances all clustered samples largely by participant (A) as compared with period relative to iron supplementation (B), indicating that the effect of iron on healthy human gut microbiotas does not overcome inter-individual variation.

**Figure S2:**
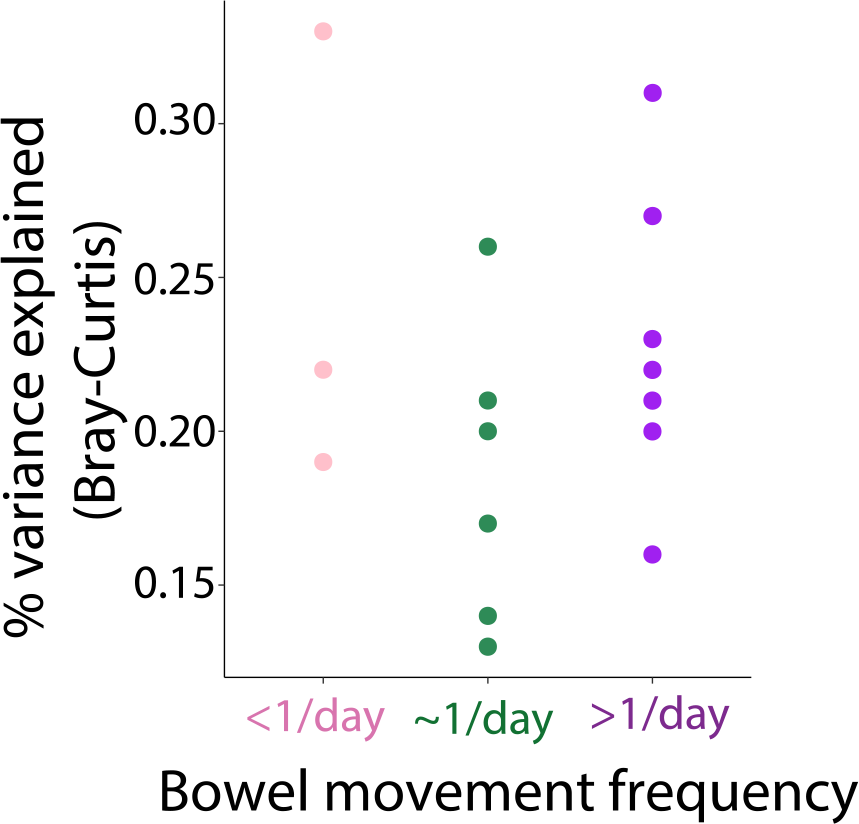
Bowel movement frequency does not explain participant-specific changes due to iron supplementation. Transit time was uncorrelated with the participant-specific community composition variance explained by iron supplementation.

**Figure S3:**
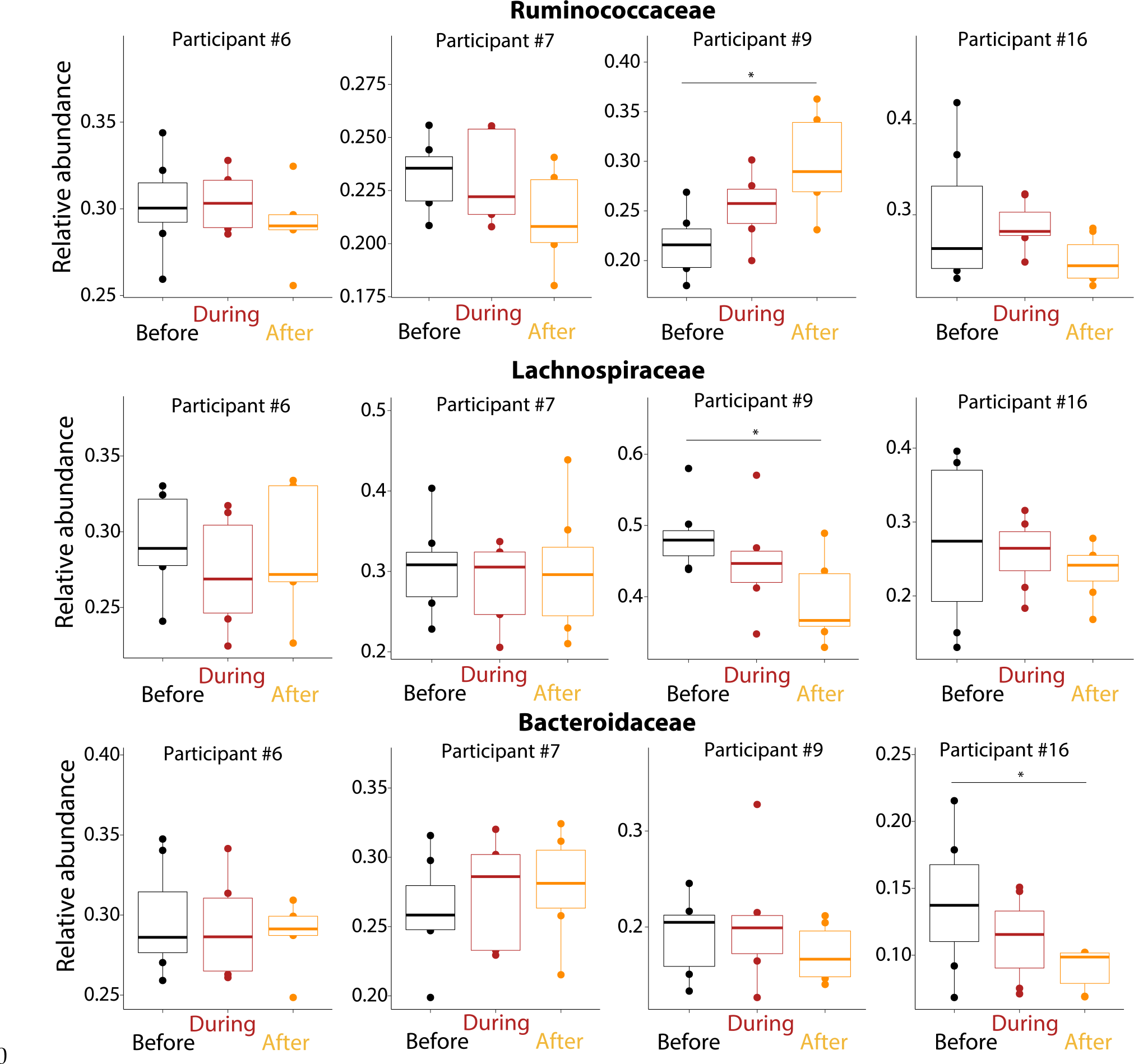
Iron supplementation does not lead to consistent changes at the family level across the most responsive participants. Across the most responsive participants (#6, #7, #9, and #16), no consistent changes were observed in the relative abundances of the Ruminococcaceae, Lachnospiraceae, and Bacteroidaceae families before, during, and after iron supplementation. *: *p*<0.05.

**Figure S4:**
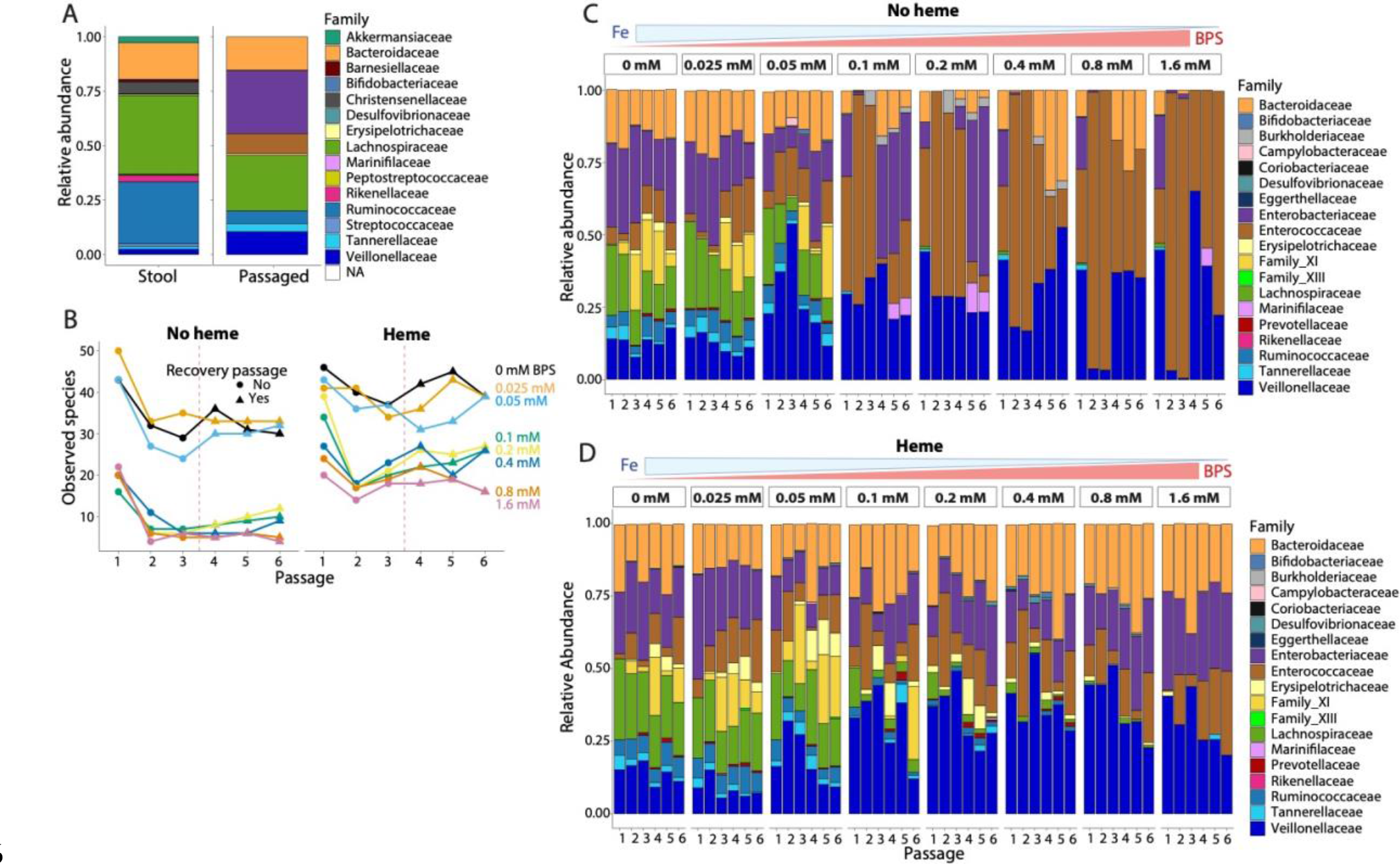
I*n vitro* communities derived from the stool of another participant undergo similar changes during and after iron deprivation as participant #16. A) An *in vitro* community derived from a pre-treatment stool sample from participant #7 retains ∼90% of the families in the original stool sample. B) Left: This community exhibited an irreversible decrease in observed species as a result of iron deprivation. Right: the decrease was ameliorated in the presence of heme. C) Participant #7’s *in vitro* community exhibited similar community changes as that of participant #16 (Figure 3): Lachnospiraceae and Ruminococcaceae members became undetectable at ≥0.1 mM BPS, Enterobacteriaceae relative abundance decreased during iron deprivation but increased during recovery up to their extinction at ≥0.4 mM BPS, and the Enterococcaceae family dominated the community at high BPS concentrations. The Burkholderiaceae family was not observed in this community. The relative abundance of the Veillonellaceae family, which was absent in participant #16’s community, increased from ∼20% to up to ∼50% during iron deprivation, suggesting that the Veillonellaceae do not depend on iron for growth and survival. D) Addition of heme rescued the same bacterial families from extinction (Enterobacteriaceae and Bacteroidaceae) as in participant #16’s community (Figure 7), and reduced the detrimental effects of iron deprivation for the Ruminococcaceae and Lachnospiraceae at ≤0.4 mM BPS, limiting the expansion of Enterococcaceae species.

**Figure S5:**
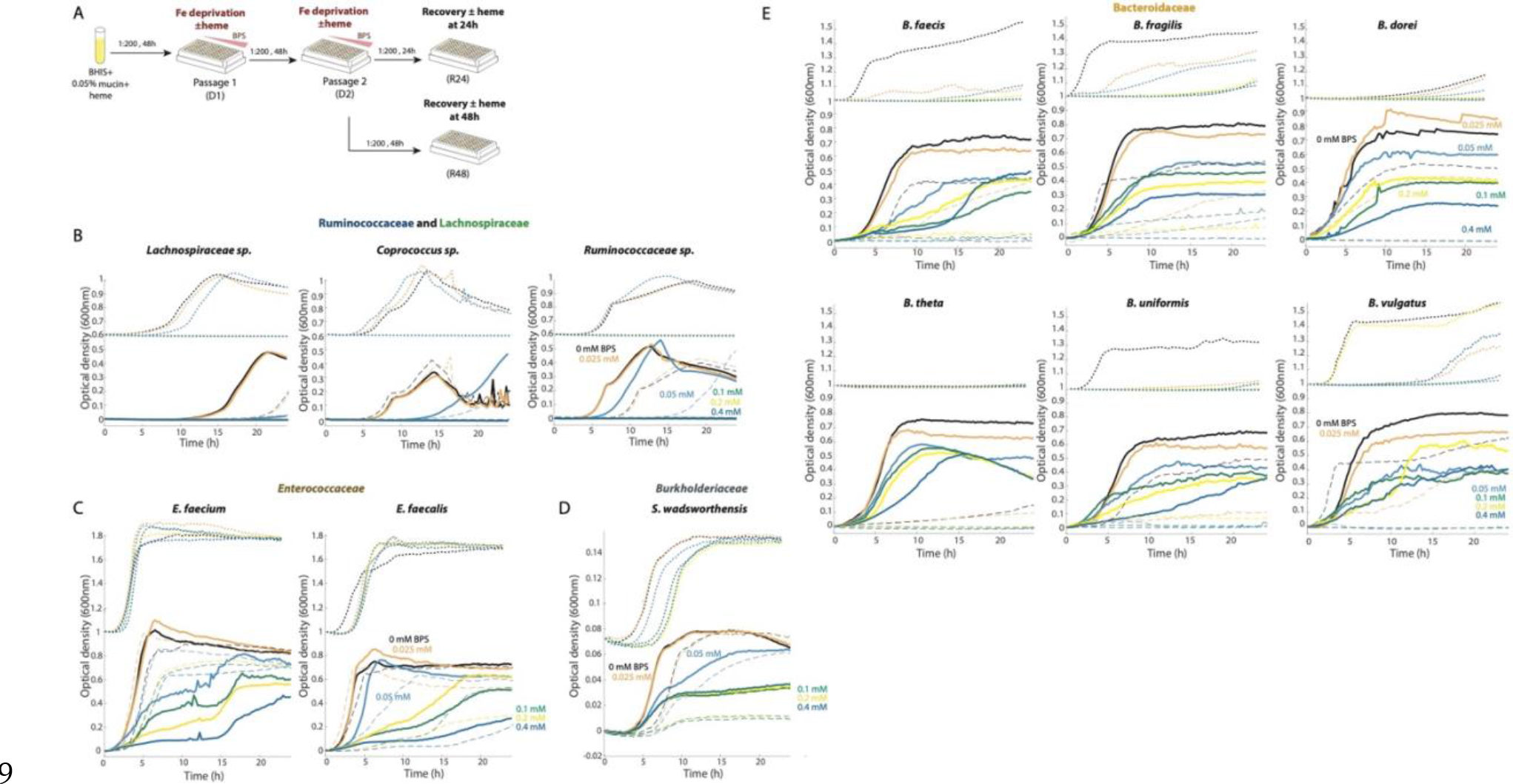
Sensitivity to iron deprivation is similar in community and isolated contexts for most species. A) Twelve isolates from five bacterial families were in grown in BHIS+0.05% mucin, with and without heme, and 0-0.4 mM BPS for two 48-h passages (iron deprivation passages D1 and D2). At the 24-h and 48-h time points of the second passage, isolates were diluted and passaged in media without BPS to access the ability to recover (recovery passages R24 and R48). B) Isolates from the Lachnospiraceae and Ruminococcaceae families exhibited species-specific growth defects at 0.05 mM BPS and failed to grow with ≥0.1 mM BPS (D1, solid lines; D2, dashed lines; R24, dotted lines). The *y*-axis for R24 curves was shifted by 0.6 for visualization purposes. C) An *E. faecium* isolate was less sensitive to iron deprivation than an *E. faecalis* isolate and exhibited more growth during a second passage in iron-deprived media than in the first. Both *E. faecium* and *E. faecalis* recovered even when treated with 0.4 mM BPS. The *y*-axis for R24 curves was shifted by 1.0 for visualization purposes. D) An *S. wadsworthensis* isolate exhibited a BPS concentration-dependent decrease in growth rate and carrying capacity, but was able to grow at all BPS concentrations. The delay in recovery was BPS concentration-dependent. The *y*- axis for R24 curves was shifted by 0.07 for visualization purposes. E) *Bacteroides* species exhibited BPS concentration-dependent and species-specific growth defects during the first iron deprivation passage. During the second iron deprivation passage, all species grew substantially worse even in the absence of BPS, illustrating the gradual effects of iron deprivation and indicating the importance of heme as a cofactor. The *y*-axis for R24 curves was shifted by 1.0 for visualization purposes.

## Notes

### Competing Interest Statement

The authors have declared no competing interest.

